# Model-based ordination for species with unequal niche widths

**DOI:** 10.1101/2020.10.05.326199

**Authors:** Bert van der Veen, Francis K.C. Hui, Knut A. Hovstad, Erik B. Solbu, Robert B. O’Hara

**Affiliations:** Department of Landscape and Biodiversity, Norwegian Institute of Bioeconomy research, Trondheim, Norway; Department of Mathematical Sciences, Norwegian University of Science and Technology, Trondheim, Norway; Centre of Biodiversity Dynamics, Norwegian University of Science and Technology, Trondheim, Norway; Research School of Finance, Actuarial Studies and Statistics, Australian National University, Canberra, Australia; The Norwegian Biodiversity Information Centre, Trondheim, Norway

**Keywords:** model-based ordination, unimodal response, niche model, unconstrained quadratic ordination, joint species distribution model

## Abstract

1. It is common practice for ecologists to examine species niches in the study of community composition. The response curve of a species in the fundamental niche is usually assumed to be quadratic. The center of a quadratic curve represents a species’ optimal environmental conditions, and the width its ability to tolerate deviations from the optimum.
2. Most multivariate methods assume species respond linearly to the environment of the niche, or with a quadratic curve that is of equal width and height for all species. However, it is widely understood that some species are generalists who tolerate deviations from their optimal environment better than others. Rare species often tolerate a smaller range of environments than more common species, corresponding to a narrow niche.
3. We propose a new method, for ordination and fitting Joint Species Distribution Models, based on Generalized Linear Mixed-Effects Models, which relaxes the assumptions of equal tolerances and equal maxima.
4. By explicitly estimating species optima, tolerances, and maxima, per ecological gradient, we can better predict change in species communities, and understand how species relate to each other.

## Introduction

One of the key topics addressed by community ecology is what causes changes in community composition. In order to explore species niches, species communities are surveyed at locations with different environmental conditions. Species tolerances are then reflected in the resulting multivariate dataset, as differences in occurrences or abundances between locations. The most favorable environmental conditions for species are represented by the optimum of the niche, where species exhibit their maximum abundance or probability of occurrence. Deviation from the optimum reflects increasingly unfavorable conditions.

Correspondence Analysis (CA) is often used to summarize community data, as it implicitly approximates the fit of a quadratic model, with the additional assumptions of equally spaced optima, sites that are well within the range of species optima, equal tolerances, and equal or independent maxima (ter Braak 1985). The combination of assuming equally spaced optima, equal maxima, and equal tolerances, gives an early niche model, called the species packing model (MacArthur & Levins 1967). The relationship of the species packing model to CA has added to its popularity among applied ecologists (Wehrden *et al*. 2009).

Recent advances in the estimation of species niches have focussed on performing ordination with explicit statistical models, such as Generalized Linear Latent Variable Models (GLLVMs; Warton *et al*. 2015). The GLLVM framework is well known for its capability to fit Joint Species Distribution Models (JSDMs; Pollock *et al*. 2014; Ovaskainen *et al*. 2017; Tobler *et al*. 2019; Zurell *et al*. 2020). In the context of JSDMs, GLLVMs assume species abundances are correlated due to similarity in response to ecological gradients, modelled with covariates or latent variables respectively. Latent variables can be understood as combinations of missing covariates, so that GLLVMs allow us to parsimoneously model species distributions. They are equivalent to ordination axes, representing complex ecological gradients (Halvorsen 2012). Recently, the use of GLLVMs to perform model-based ordination has increased in popularity (Inoue *et al*. 2017; Björk *et al*. 2018; Lacoste *et al*. 2019; Damgaard *et al*. 2020).

With intercepts included for row standardization, GLLVMs fit the species packing model (Jamil & ter Braak 2013; Hui *et al*. 2015), though with maxima that are equal for the latent variables. Existing GLLVMs assume that latent variables are linear, just as all classical ordination methods (Jamil & ter Braak 2013). However, it is widely understood that species have unequal tolerances and maxima, so that the assumptions of linear latent variables, and equal tolerances, are unlikely to hold in practice.

In this paper, our goal is to overcome the assumptions of equal tolerances, and equal maxima, by formulating a GLLVM with quadratic latent variables. To our knowledge, there has been no attempt to implement such a GLLVM until now. Although seemingly a straightforward extension, the quadratic term explicitly allows species niches to be estimated without constraints on the parameters. This means that optima, tolerances, and maxima per latent variable, as well as the lengths of ecological gradients, can all be explicitly estimated. Explicitly estimating the combination of these three parameters gives unique insight into reasons for species low detectability, whether it is due to low abundance or probability of occurrence (maxima), a high degree of habitat specialization (tolerance), or due to unsuitable observed environmental conditions (optima). In combination with knowledge of the study system, these parameters can help ecologists to determine why certain species are rare. Additionally, due to the model-based nature of the proposed ordination method, it is possible to calculate confidence intervals for each set of parameters, providing unparalleled benefits for inference when using ordination. In the context of JSDMs, the quadratic GLLVM models latent species distributions, without covariates in the model. When covariates are included, the quadratic GLLVM partitions species distributions in observed (fixed effects) and latent or unobserved (random effects), similar to the partitioning of fixed and random effects in mixed-effects models when covariates are included.

In contrast to classical ordination methods, GLLVMs model the latent variables as unobserved, treating them as random rather than fixed (Walker & Jackson 2011), which consequently have to be integrated over in the likelihood. Here, we develop a variational approximations (VA) implementation after Hui *et al*. (2017) and Niku *et al*. (2019a), to perform calculations quickly and efficiently. In addition to presenting the quadratic GLLVM, we perform simulations to evaluate the accuracy of the VA implementation, and the capability of the quadratic GLLVM to retrieve the true species-specific parameters and latent variables. We use two real world datasets to demonstrate use and interpretation of the proposed quadratic GLLVM: 1) a small dataset of hunting spiders in a Dutch dune ecosystem (van der Aart & Smeek-Enserink 1974), and 2) a larger dataset on Swiss alpine plant species on a strong elevation gradient (D’Amen *et al*. 2018).

## Model formulation

The ecological niche is here described by a quadratic function involving three parameters; the optimum ***u***_*j*_, the tolerance ***t***_*j*_, and the maximum ***c***_*j*_. The optimum ***u***_*j*_ is the location on the ecological gradient where a species exhibits its highest abundance or probability of occurrence (the maximum ***c***_*j*_). The tolerance ***t***_*j*_ is a measure of the width or breadth of the niche, and indicates if a species is a generalist or specialist.

Consider an *n* × *p* matrix of observations, where *y*_*ij*_ denotes the response of species *j* = 1 *… p* at site *i* = 1 *… n*. Then in the quadratic GLLVM, we assume that, conditional on a vector ***z***_*i*_ of *q* = 1 *… d* latent variables where *d* ≪ *p*, the responses *y*_*ij*_ at site *i* are independent observations from a distribution whose mean, denoted here as E(*y*_*ij*_|***z***_*i*_), is modelled as:

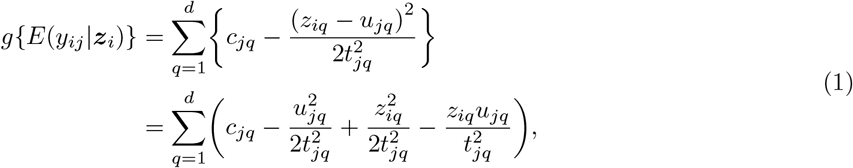

where *g*{·} is a known link function (e.g. the log-link when the responses are assumed to be Poisson, negative-binomial, or gamma distributed, the probit-link when the responses are assumed to be Bernoulli or ordinal distributed, and the identity-link for responses that are assumed to be Gaussian distributed).

To facilitate easier estimation, and for a closer comparison to the linear GLLVM, we formulate the quadratic GLLVM in matrix notation:

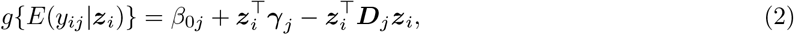

with a species-specific intercept *β*_0*j*_ that accounts for e.g. mean abundances, and a vector of coefficients per species for the linear term ***γ***_*j*_. We can see a third term is added here to the existing structure of the linear GLLVM, which models tolerances and maxima per species and latent variable. Specifically, we introduce a diagonal matrix ***D***_*j*_ of quadratic coefficients with each diagonal element being the quadratic effect for latent variable *q* and species *j*. We require ***D***_*j*_ to be a positive-definite diagonal matrix, to ensure concave curves to the latent variables. Thus, 2***D***_*j*_ is the precision matrix of the ecological niche (likewise (2***D***_*j*_)^*−*1^ is the covariance matrix). Additionally, row intercepts or covariates can be included as in Hui *et al*. (2017), or species traits as in Niku *et al*. (2019a), though we have chosen to omit those terms here and focus on the case of unconstrained ordination.

With ***D***_*j*_ being a diagonal matrix with the positive elements *D*_*jqq*_, the vector of species maxima ***c***_*j*_ with elements *c*_*jq*_, the vector of species optima ***u***_*j*_ with elements *u*_*jq*_, and the vector of species tolerances ***t***_*j*_ with elements *t*_*jq*_, we derive the following connections between the parameters in equations (1) and (2): 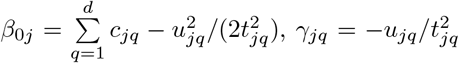, and 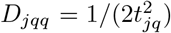. Similarly, for the formulation in equation (2), the parameters in equation (1) can be retrieved: 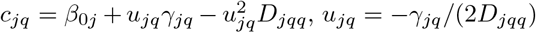, and 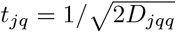.

Four special cases of the quadratic GLLVM, as formulated in equation (2), are worth discussing: 1) ***D***_*j*_ = ***D***, i.e. common tolerances for species, 2) ***D***_*j*_ = *D*_11_**I**_*d*_ where **I**_*d*_ is a *d* × *d* identity matrix, i.e. equal tolerances for species and latent variables, 3) when ***D***_*j*_ = 0 for a subset of the *p* species, and 4) when ***D***_*j*_ = 0 for all *p* species. The first case assumes tolerances to be the same across species, but not latent variables, and additionally places constraints on the species maxima. This species-common tolerances model might prove useful in practice, as it requires fewer observations per species than the full quadratic GLLVM, but still explicitly includes quadratic latent variables. The second case can be shown to be equivalent to the linear GLLVM with row intercepts as presented in Hui *et al*. (2015), which assumes tolerances to be the same for all species and latent variables, and the maxima to be the same for all latent variables. In the third case, some species respond to the latent variable linearly, while others exhibit quadratic responses. The fourth case is the most basic GLLVM with linear latent variables, currently possible to fit with e.g. `boral` (Hui 2016), `HMSC-R` (Tikhonov *et al*. 2020), and `gllvm` (Niku *et al*. 2020).

## Model interpretation

In this section, we derive and discuss various tools that are commonly used in the application of JSDMs and ordination, such as calculating residual correlations and partitioning residual variance, calculating gradient length, and visualizing the ordination, and demonstrate how they can be adapted to the proposed quadratic GLLVM.

### Residual covariance matrix

One aspect GLLVMs are known for is modelling species residual correlations, calculated from the residual covariance matrix (Zurell *et al*. 2018; Blanchet *et al*. 2020). To facilitate calculation of the residual covariance matrix, we can reparameterize all GLLVMs as a multivariate mixed-effects model with a residual term:

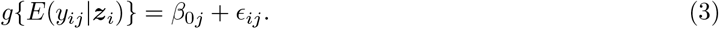

Here, *ϵ*_*ij*_ accounts for any residual information that is not accounted for by fixed-effects in the model, such as covariates or intercepts (Warton *et al*. 2015). Assuming the latent variables are independent for all sites, the elements of the residual covariance matrix are given by:

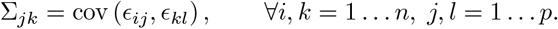

For a a length *p* vector *ϵ*_*i*_, existing JSDM implementations assume *ϵ*_*i*_ ∼ 𝒩 (**0, Σ**), i.e. the residual term follows a multivariate normal distribution. For the linear GLLVM, it is straightforward to show that 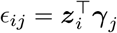, so it follows that 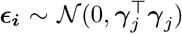. In essence, GLLVMs peform a low rank approximation to the covariance matrix of a residual term. The rank of this residual covariance matrix is equal to the number of estimated latent variables *d* in the model for the linear GLLVM.

Turning to the quadratic GLLVM, where 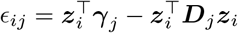, the elements of the residual covariance matrix are:

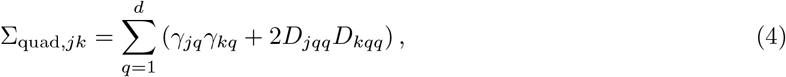

for which a proof is given in Appendix S1. This can be rewritten in terms of the species optima ***u***_*j*_ and tolerances ***t***_*j*_:

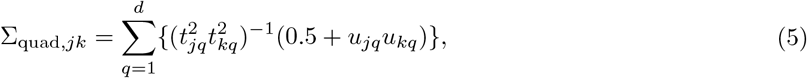

from which it follows that 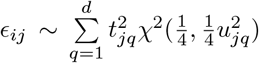, in words: the residual term follows a generalized χ^2^ distribution (Khatri 1980).

Equation (4) and equation (5) additionally serve to demonstrate how to partition the residual variance of the quadratic GLLVM, e.g. per latent variable, for the linear and quadratic term separately, or both. Variance partitioning is commonly used in the application of ordination methods, e.g. to determine fit (Økland 1999), or to explore causes of residual variance (Borcard *et al*. 1992; Økland & Eilertsen 1994). Covariates can be included in the model to account for the residual variance otherwise accounted for by the latent variables. The residual variance can be used to identify indicator species i.e. those species that best represent an ecological gradient, or to calculate a measure of *R*^2^ (Nakagawa & Schielzeth 2013).

The rank of the residual covariance matrix is double that of a linear GLLVM with the same number of latent variables: 2*d*. The additional quadratic term thus allows us to account for more residual correlations between species, with fewer latent variables. This corresponds with the ecological notion that species often respond to few major complex ecological gradients (Halvorsen 2012). From this, we see that when the number of latent variables in a quadratic GLLVM exceeds 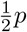, there are more parameters included than in a JSDM with an unstructured residual covariance matrix. However, this is not an issue here, since for ordination purposes we are only interested in cases where there are much fewer latent variables *d* than species *p*.

### Gradient length

The length of an ecological gradient is of great interest to ecologists in the use of ordination, because it provides a measure of beta diversity (Oksanen & Tonteri 1995). Longer gradients indicate higher diversity, as spacing (i.e. dissimilarity in the species community) between sites in latent space is potentially larger. In the past, it has been emphasized that short gradients are better analysed using linear ordination methods, and longer with unimodal methods (ter Braak & Prentice 1988). However, the quadratic GLLVM allows species to exhibit both linear and unimodal responses, and so it is appropriate for both, and it is no longer required to switch ordination method as a consequence of gradient length.

To determine gradient length from the proposed quadratic GLLVM, we define the ecological gradients 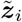, as a function of the latent variables ***z***_*i*_, but with a diagonal covariance matrix ***G*** of size *d* × *d*. First, for a species-common tolerances model, we note that the quadratic term in equation (2), i.e. 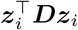, can instead be written as 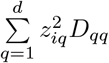, so that 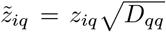, and 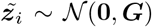, where *G* = *D*. Then, the gradient length is approximately 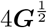 (i.e. the approximate width of a normal distribution), and ***D*** = **I**

For the species-specific tolerances model, we note that one of the uses of gradient length in the past has been to rescale the latent variables so that an ordination diagram can be understood in terms of compositional turnover (Hill & Gauch 1980). This requires the mean species tolerances to be one (as is the case for the species-common tolerances model above), so that the covariance matrix of the ecological gradient in the species-specific tolerances model is 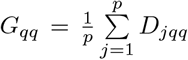, so the matrix of quadratic coefficients ***D***_*j*_ is scaled by the inverse of the covariance matrix of the ecological gradient, ***G***^*−*1^. However, we choose to use the median of the species tolerances instead, as it more accurately represents gradient length with both linear and quadratic responses of species in the model. In general, the proposed quadratic model allows further exploration of measures of gradient length by, for example, using the mean tolerance of species with clear quadratic responses, rather than the median of all tolerances.

The measure of gradient length calculated here, can be interpreted in the same manner as the gradient length provided by Detrended Correspondence Analysis (Hill & Gauch 1980).

### Ordination diagram

Usually, a biplot (Gabriel 1971) is constructed to visually inspect results from an ordination. For the quadratic GLLVM, biplots tend to create an arch when the residual variance of the linear term is smaller than the residual variance of the quadratic term.

Instead, we propose that species optima and tolerances can be plotted directly, so that species niches are visualized in a two-dimensional latent space from a top-down perspective. The widths of the niches can then potentially be represented as ellipses using the estimated species tolerances (i.e. providing species distributions in latent space), so that co-occurrence patterns can be inferred from the (lack of) overlap between ellipses. Additionally, information on sites, such as the predicted locations and prediction regions, can be added (Hui *et al*. 2017). Information for the sites can be used to infer the distance of sites to the species optima (i.e. the suitability of sites for species), or to the egdes of species niches (see the hunting spiders example below).

Finally, based on the disscussion in the two subsections above, there are two ways of scaling the ordination diagram: 1) by the residual variance per latent variable, or 2) by the mean or median tolerance. In the first scaling, the diagram is scaled to draw attention to the latent variable that explains most variance in the model. However, the second scaling has a more ecological intuitive interpretation. If the tolerances are assumed to be common for species, the second scaling provides an ordination diagram in units of compositional turnover (Gauch 1982). When the linear and quadratic terms in the model explain an equal proportion of the total residual variance per latent variable, these scalings produce similar results.

## Model estimation

We propose to use variational approximations (VA; Hui *et al*. 2017) for estimation and inference for the quadratic GLLVM. Broadly speaking, VA is a general technique used to provide a closed-form approximation to the marginal log-likelihood of a model with random effects or latent variables, when an analytical solution is not available. Computationally, VA can be orders of magnitude faster than MCMC, numerical integration, or even the Laplace approximation (Niku *et al*. 2019a), and without loss of accuracy (Hui *et al*. 2017). However, the calculation of the VA log-likelihoods needs to be derived on a case-by-case basis. In contrast, the Laplace approximation can be applied automatically in many cases (Kristensen *et al*. 2016). Note it is not possible to approximate the marginal likelihood of a quadratic GLLVM with the Laplace approximation (K. Kristensen, pers. comm., March 8th 2019).

The marginal log-likelihood of a quadratic GLLVM is given by:

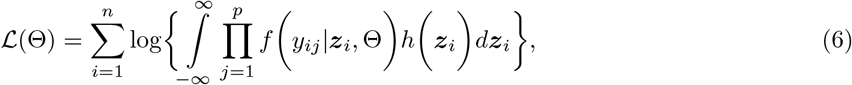

where *f* (*y*_*ij*_|***z***_*i*_, Θ) is the distribution of the species responses given the latent variables. As mentioned previously, and as per Hui *et al*. (2015), we assume the distribution of the latent variables *h*(***z***_*i*_) to be multivariate standard normal i.e. *h*(***z***_*i*_) = 𝒩(**0, I**). The vector Θ includes all parameters in the model Θ = {*β*_01_ *… β*_0*j*_, *γ*_11_ *… γ*_*jq*_, *D*_111_ *… D*_*jqq*_}^*T*^. Equation (6) can be straightforwardly modified if covariates are also included in the quadratic GLLVM.

In VA, we construct a lower bound to equation (6), by assuming that the posterior distribution of the latent variables can be approximated by a closed form distribution e.g., a multivariate normal distribution. We then minimze the Kullback-Leibler divergence between this approximate closed-form distribution (also known as the variational distribution) and the true posterior distribution. Hui *et al*. (2017) showed that, for GLLVMs with linear latent variables, the optimal variational distribution is multivariate normal ***z***_*i*_ ∼ 𝒩 (***a***_*i*_, ***A***_*i*_), with mean ***a***_*i*_ and covariance matrix ***A***_*i*_, so we will adopt this choice here as well.

In Appendix S1 we provide information on calculating approximate confidence intervals for the parameters. In Appendix S2 we provide derivations for the log-likelihood of common response types in community ecology, such as count data (Poisson, and negative-binomial with quadratic mean-variance relationship, and both assuming a log-link function), binary data and ordinal data (both with probit-link function), as well as positive continuous data (gamma, with log-link function) and continuous data (Gaussian, with an identity-link function).

### Simulation study

To assess how well the proposed model retrieves the true latent variables ***z***_*i*_, optima ***u***_*j*_, tolerances ***t***_*j*_, and maxima ***c***_*j*_, we performed simulations for six response distributions; 1) Gaussian, 2) gamma, 3) Poisson, 4) negative-binomial, 5) Bernoulli, and 6) ordinal. The R-code used for the simulations is provided in Appendix S3. For each of the distributions, we simulated 1000 datasets with different numbers of sites and species. A consequence of the negative-only third term in the quadratic GLLVM, is that the model often simulates a large number of zeros (more so than the linear GLLVM), providing a challenge in testing its accuracy, especially for small datasets. First, to study the accuracy of the VA approximation, we simulated datasets of *p* = 20 to 100 species in increments of 10, while keeping the number of sites constant at *n* = 100. Hui *et al*. (2017) argued that the VA log-likelihood is expected to converge to the true likelihood as *p* → ∞ (i.e. for a large number of species), thus this will allow us to study the finite sample properties of the VA approximation for the proposed model.

Second, to explore the sample size required to accurately estimate the species-specific parameters e.g., species optima ***u***_*j*_, tolerances ***t***_*j*_, and maxima ***c***_*j*_, we simulated datasets of *n* = 20 to 100 sites in increments of 10, while keeping the number of species constant at *p* = 100. For each dataset, we compared 12 combinations of initial values and fitting algorithms (see Appendix S4: Fitting, for details), and picked the model with the highest log-likelihood (see Appendix S5: Fig. S1 for the frequency at which different types of initial values and fitting algorithm were used in the best models per distribution).

As a true model, we considered a quadratic GLLVM with *d* = 2 latent variables, which was constructed as follows. First, the species-specific intercepts *β*_0*j*_ were simulated as Uniform(−1, 1), which corresponds to species with low abundance or occurrence. Next, the true coefficients corresponding to the linear terms in the model ***γ***_*j*_, were simulated independently as Uniform(−5, 5), and the true quadratic coefficients as Uniform(−5, −0.5). The true latent variables were simulated as ***z***_*i*_ ∼ 𝒩 (**0, I**). For the Gaussian, negative-binomial, and gamma distribution, the dispersion parameter for all species was set equal to one. For the ordinal distribution we assumed six classes with the true cut-offs being 0, 1, 2, 3, 4, 5, meaning that species were most often absent (category 1), while they were rarely very abundant (category 6).

We measured performance of the quadratic GLLVM by the prediction of the latent variables ***z***_*i*_ and the species optima ***u***_*j*_. The species optima are a function of both the linear and quadratic coefficients and should provide a good overall measure of performance for retrieving the true species-specific parameters, in addition to being of specific interest to ecologists. Though it is common to measure the performance of ordination methods using the Procrustes error (Peres-Neto & Jackson 2001), we chose to use the Median Absolute Error (MAE) instead, as we often observed a highly skewed error distribution for the species optima. Additionally, interpreting the MAE is more intuitive, as it measures the deviation from the truth in the same units as the coefficients of interest. We excluded the first optimum of the second latent variable as this was fixed to zero for reasons of parameter identifiability (Hui *et al*. 2015), and excluded optima that could not be estimated. Since the quadratic GLLVM allows species to exhibit linear responses, which have infinite optima, we chose to remove all optima larger than 10 and smaller than −10, i.e. for those species that lacked a sufficiently strong quadratic signal in the simulated datasets. Including these optima would result in a biased view of the accuracy of the optima that can be estimated by the model. This process resulted in a vector of optima, which we then used to calculate the MAE. For clarity and transparency, we additionally present the number of optima removed for each of the datasets, to further provide an impression of the data requirements of the proposed quadratic GLLVM.

For all of the models fitted to Gaussian and gamma response datasets, typically none or only a few optima were excluded, meaning that the median number excluded was zero. In general, and not surprisingly, more optima were excluded for models fitted to datasets where *n/p* was small and for discrete distributions. For example, when *n* = 20 sites and *p* = 100 species, the median number of optima excluded for datasets with Poisson responses was 4 (2 – 8, first and third quartiles), for datasets with negative-binomial responses this was 7 (5 – 10), for datasets with Bernoulli responses this was 31 (24 – 38), and for datasets with ordinal responses this was 17 (13 – 23). In contrast, for datasets where *n/p* was large, considerably less optima were excluded across all response types. For example, when *n* = 100 and *p* = 100, for Poisson and negative-binomial response datasets the median number of excluded optima was zero, while for Bernoulli response datasets the median number of optima excluded was 4 (2 – 5), and for ordinal response datasets this was 2 (1 – 3).

The MAE per distribution and for the different sized datasets is presented in Figure 1 (see Appendix S5: Fig. S2 for the same figure with all species optima). As expected, the quadratic GLLVM was more accurate for datasets with larger *p* and larger *n*. For all distributions, the latent variables were often better retrieved than the species optima. This is not surprising, as the species optima are a function of two parameters, particularly the inverse of the quadratic coefficients, so that a small change in the quadratic coefficients can result in a large change in the species optima. When fitted to Gaussian or gamma response datasets, regardless of the dimensions of the data, the model performed best. The accuracy of the estimated species optima was only slightly lower for the Poisson distributed datasets with 70 or more sites, while the latent variables were accurately estimated even with small *p*. These results are consistent with the results above regarding the number of excluded optima. Although the accuracy of species optima for negative-binomial response datasets seems similar to that of Poisson response datasets, this is inconsistent with the number of excluded optima reported above, as that was considerably larger for the negative-binomial. The model was not accurate for Bernoulli or ordinal response datasets with small *n* and *p*. However, when the number of sites and species increased above 40, the performance of the quadratic GLLVM in these cases improved considerably, consistent with the number of excluded species optima.

**Figure 1:**
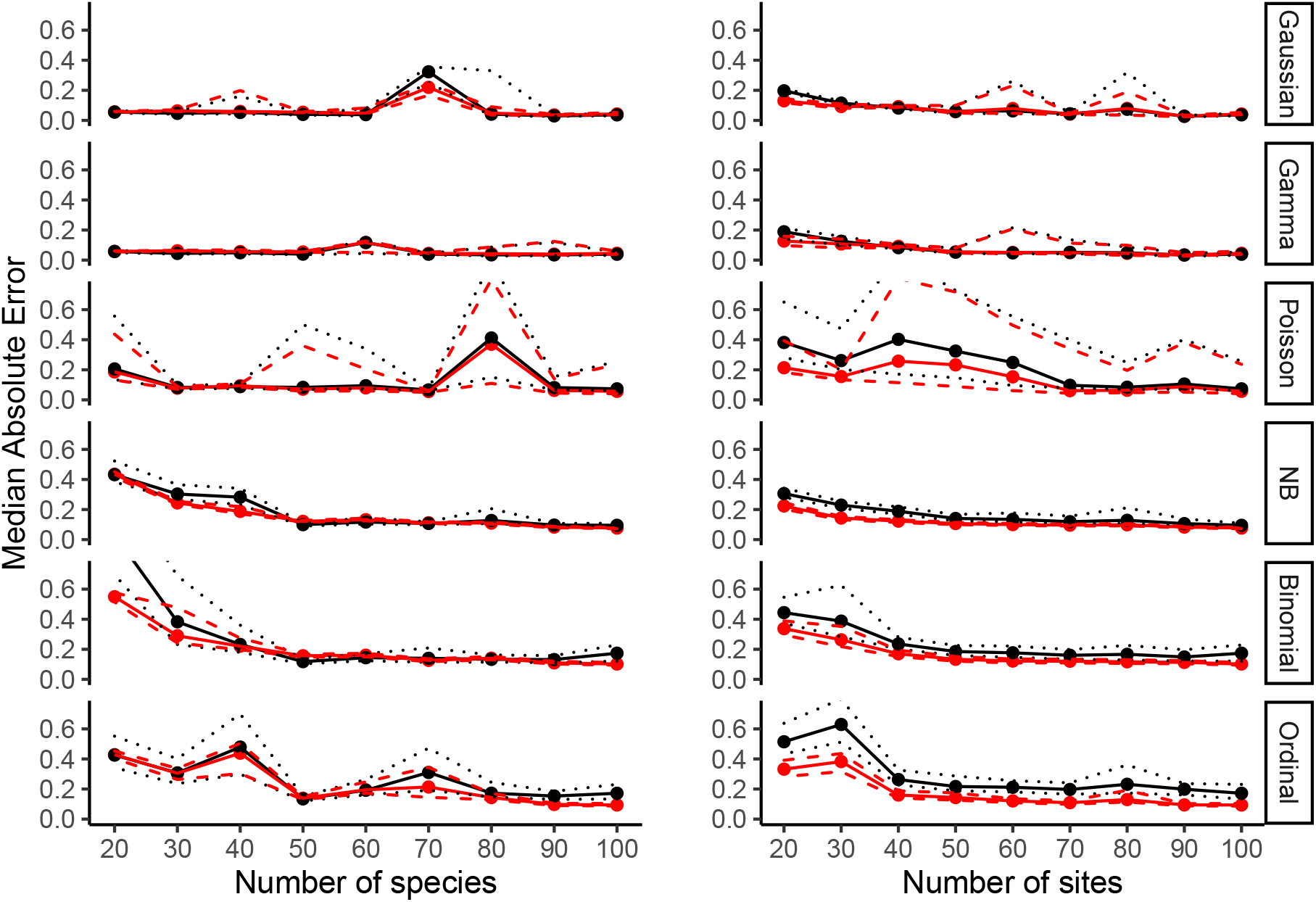
Simulation results for the 1000 best fitting quadratic GLLVMs across six different response distributions, with the MAE calculated based on optima that could be estimated (optima outside the range (−10,10) were excluded). The left column shows simulations where the number of sites was kept constant at n = 100, and analogous for the right column with p = 100. The figure includes the median MAE for species optima (black) and latent variables (red), with the first and third quartiles represented as dotted (optima) and dashed (latent variables) lines.

## Applications to real data

We applied the proposed quadratic GLLVM to two different datasets: 1) the classical hunting spiders dataset collected by van der Aart & Smeek-Enserink (1974) in Dutch dunes, available in the `mvabund R` package (Wang *et al*. 2012), and 2) a dataset of plants in the Swiss Alps (available in the dryad database; D’Amen *et al*. 2017).

### Hunting spiders

For the hunting spiders dataset, van der Aart & Smeek-Enserink (1974) used pitfall traps to collect spiders over a 60 week period, resulting in a dataset of counts for each of the *n* = 28 sites and *p* = 12 species. It has been used in the testing of ordination methods before (e.g. ter Braak 1985, 1986; Yee 2004; Hui *et al*. 2015), providing some reference results for comparison here. To find the model that best fitted the hunting spiders dataset, and to limit the number of required model fits, we first performed model selection using Akaike’s Information Criterion (AIC; Burnham & Anderson 2002) on linear GLLVMs with *d* = 1 to 5 latent variables, and with Poisson distributions. Though the hunting spiders dataset exhibits overdispersion in the linear GLLVM (Hui *et al*. 2015), the quadratic GLLVM models overdispersion with the latent variables (see Appendix S2: Negative-Binomial: overdispersed counted responses). Second, we fitted a linear GLLVM with random row intercepts (i.e. equal tolerances), a quadratic GLLVM with common tolerances, and a quadratic GLLVM with unequal tolerances, to determine which model structure was most suited for the data. Third, with the best model structure from step two, we again tested for the optimal number of latent variables, after which we explored different sets of initial values and fitting algorithms to find the model that maxmizes the VA log-likelihood (see Appendix S5 for the results).

The best model of step one included *d* = 2 latent variables, the best model from step two included unequal tolerances, and the best model from step three included *d* = 3 latent variables. The results for te first two latent variables of the final model fit, which explained most residual variation, are presented in Figure 2. We used the residual variance to determine which latent variables explained most variation i.e. were most important to consider for inference. For the quadratic GLLVM, the first and second latent variables explained most variation in the model; 40% and 57% respectively. Overall, the quadratic GLLVM explained four and a half times more residual variation than a linear GLLVM with the same number of latent variables. The lengths of the first two ecological gradients were 5.46 (4.28-6.64, 95% confidence interval), and 3.35 (3.14- 3.55). The confidence interval of the gradient length for the third ecological gradient included zero, so we do not present results of that here.

**Figure 2:**
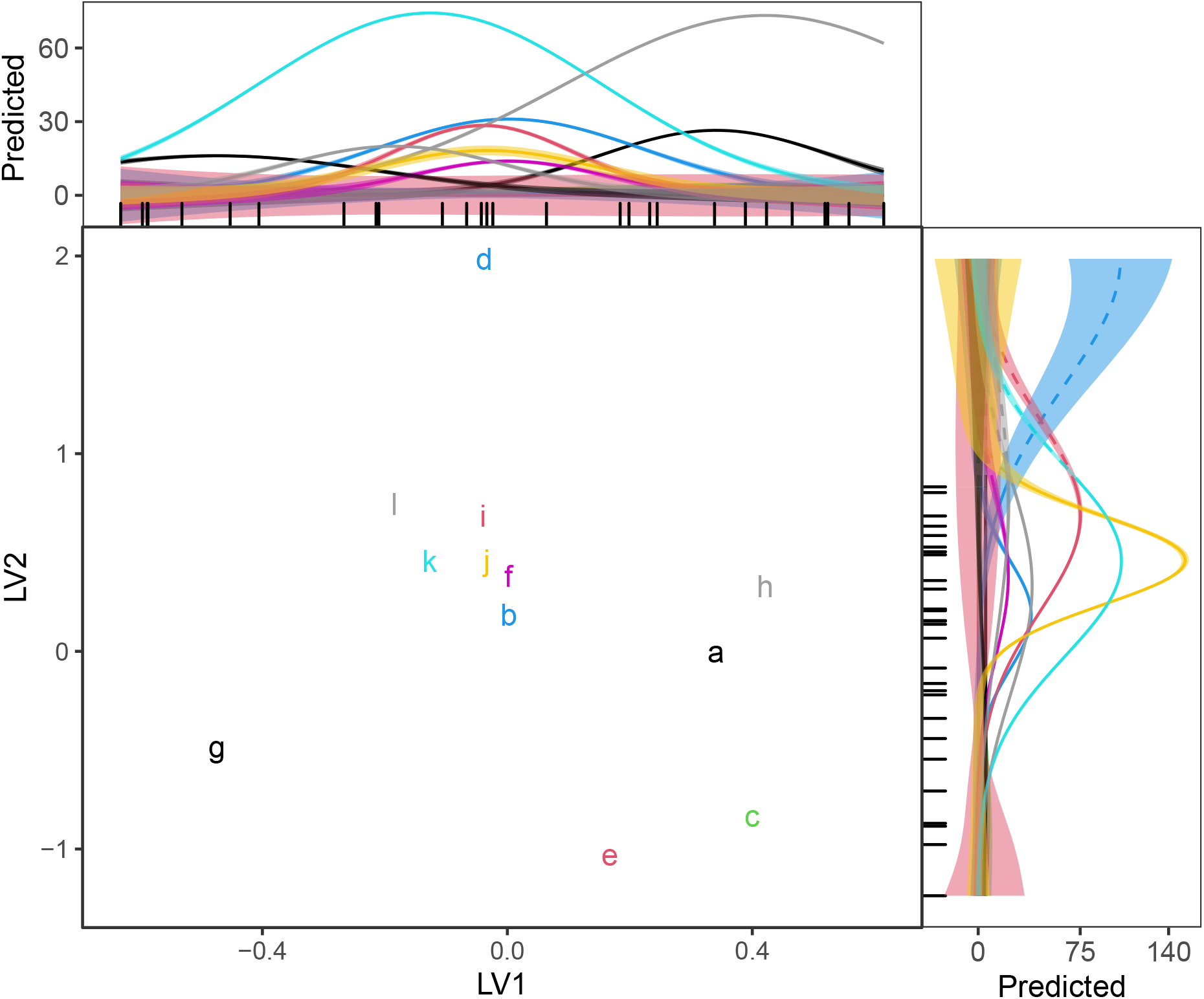
Ordination plot for the first two latent variables of the final quadratic GLLVM fit to the hunting spiders dataset, scaled by the residual variances. Species optima are shown as letters, indicating the follwing species: a = *Alopecosa accentuata*, b = *Alopecosa cuneata*, c = *Alopecosa fabrilis*, d = *Arctosa lutetiana*, e = *Arctosa perita*, f = *Alonia albimana*, g = *Pardosa lugubris*, h = *Pardosa monticola*, i = *Pardosa nigriceps*, j = *Pardosa pullata*, k = *Trochosa terricola*, l = *Zora spinimana*. Species quadratic curves are included as side panels, with dashed lines indicating unobserved parts of species niches, and with bands representing 95% confidence intervals. Site locations and prediction regions have not been included, in favor of readability.

Ter Braak (1985) and Yee (2004) both visualized quadratic curves of the first latent variable using variations of Poisson regression and generalized additive models, respectively. There are clear similarities between the species response curves for the first latent variable in Figure 2, and the corresponding response curves described by ter Braak (1985) and Yee (2004). Though ter Braak (1985) concluded all species exhibited unimodal curves on the first latent variable (without formal testing), the species *Alopecosa fabrilis, Arctosa perita* and *Pardosa lugubris* had confidence intervals, for the quadratic coefficients, that include zero in the quadratic GLLVM. For the following conclusions, species with confidence intervals of quadratic coefficients that crossed zero were excluded.

Turning to the species niches, on the first latent variable all optima were observed (i.e. within the range of the latent variable), and on the second latent variable only the optimum of *Arctosa lutetiana* was unobserved. On the first latent variable, *Aulonia albimana* had the lowest maximum and *Trochosa terricola* the highest. On the second latent variable, *Alopecosa accentuata* had the lowest maximum, and *Pardosa pullata* the highest. On the first latent variable it was possible to distinguish that *Trochosa terricola* and *Pardosa monticola* had wider niches than *Pardosa nigriceps, Aulonia albimana*, and *Arctosa lutetiana*, and the confidence intervals of these two groups did not overlap. On the second latent variable, *Pardosa pullata* had the most narrow niche, and *Pardosa monticola* the widest, and the confidence intervals of the tolerances for these species did not overlap. *Alopecosa cuneata* was more tolerant to changes in the environment than *Pardosa pullata* but less than *Pardosa monticola*. Additionally, *Alopecosa fabrilis, Trochosa terricola*, and *Pardosa nigriceps* were more tolerant to changes in the environment than *Pardosa pullata*, though it was not possible to say if this was more or less than *Pardosa monticola*. Overall, *Arctosa lutetiana* had the smallest tolerance across all three latent variables.

Overall, due to a combination of low maxima and low tolerances, *Arctosa lutetiana* is predicted to be most prone to changes in the environment of the first latent variable, and for the second latent variable *Arctosa perita*.

### Swiss alpine plants

In the second application, *n* = 912 plots of 4 *m*^2^ each were used to record binary data on *p* = 175 plant species. Plots were located on a strong elevation gradient ranging from 375 meters to 3210 meters above sea level (D’Amen *et al*. 2018). We excluded 72 plots without any presences, and 103 plots with less than six presences, though it is possible to run the model including these plots, so that the final dataset included *n* = 737 plots. Species with less than 20 presences were excluded by the original study (D’Amen *et al*. 2018), though it would not have presented a problem for the quadratic GLLVM had we included those here. Instead of selecting the optimal number of latent variables, we directly fitted the model to the data, using the Bernoulli distribution and with *d* = 2 latent variables, for the purpose of creating an ordination diagram. We tested different sets of initial values and retained the model that had the highest log-likelihood.

The first latent variable explained 83% of the overall residual variation in the model, of which 61% was accounted for by the linear term. The length of the first ecological gradient was 3.52 (2.85-4.18, 95% confidence interval). Since the first latent variable explained considerably more residual variation than the second, we here focus our inference on that alone for illustration purposes. The species response curves for the first latent variable are visualized in Figure 3a-c. To improve readability, species are numbered by their location in the dataset, for which the corresponding names are included in Figure 4. In Figure 4 species tolerances for the first latent variable are visualized, with approximate 95% confidence intervals.

**Figure 3:**
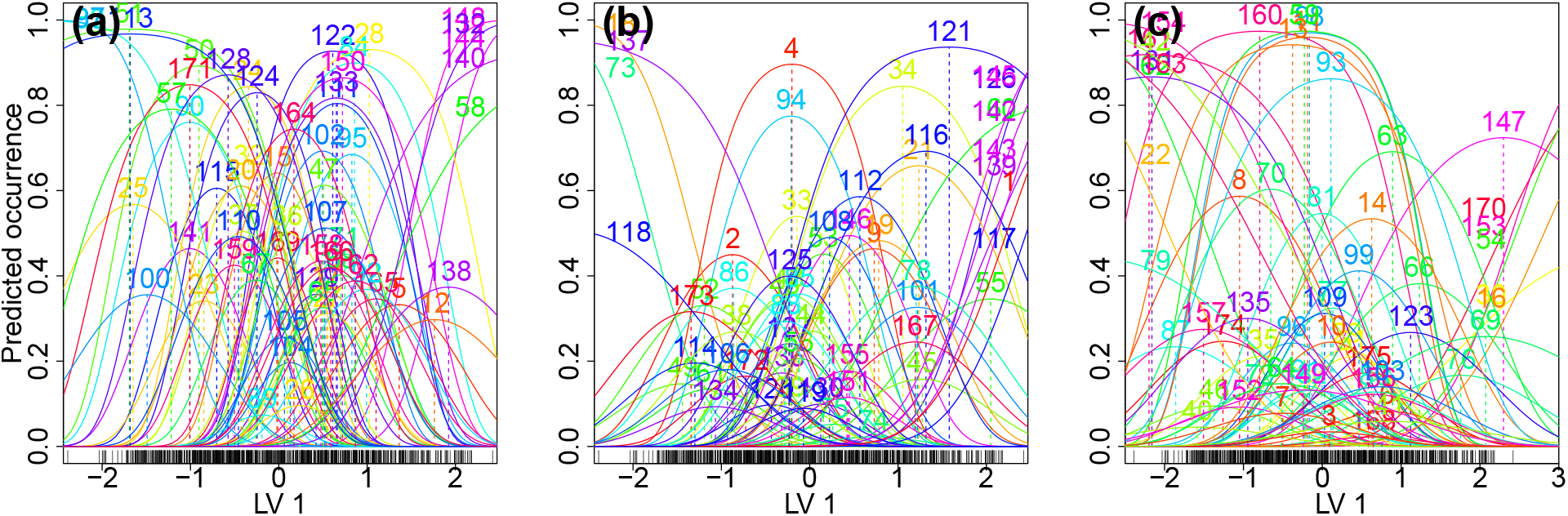
One-dimensional figures for the quadratic GLLVM fit to the Swiss alpine plants dataset. Each plot includes approximately one third of the species in the dataset, which have been sorted based on their variation explained, so that the first plot includes species explaining most of the variation. Plot a) represents 56% of the residual variation, plot b) represents 28% of the residual variation, and plot 2) represents 16% of the residual variation. Dashed coloured lines indicate the position of species optima, and the rug plot at the bottom indicates predicted locations of the plots. The numbers correspond with the species names in Figure 4.

**Figure 4:**
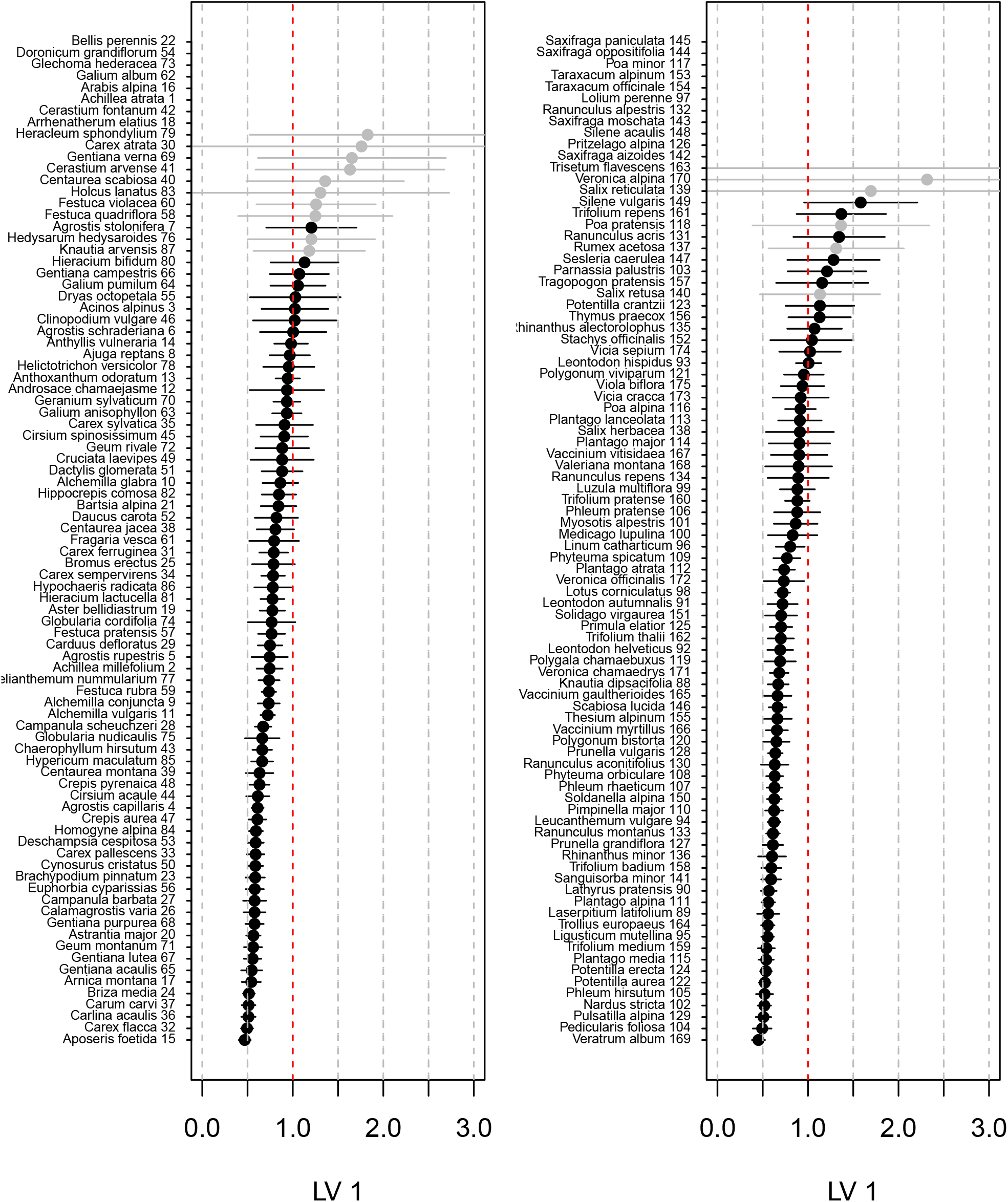
Species tolerances and approximate 95% confidence intervals derived using the Delta method, for the Swiss plants dataset. When optima are outside the range of the latent variable, or when tolerances cross one (indicated with a red dashed line), species have partially unobserved niches. The panels show the first and second half of species in the dataset respectively, ordered by the size of their tolerances. Species for which the confidence interval for the quadratic coefficients crosses zero are shown in grey. Species at the top of the plot, seemingly without tolerances, exhibit near linear responses, so that their tolerances are very large. Grey dashed lines are added at increments of 0.5 as visual aid.

Environmental tolerances from species of which the confidence interval for the quadratic coefficients on the first latent variable did not include zero, ranged from 0.45 (*Veratrum album*) to 1.58 (*Silene vulgaris*) with a median tolerance of 0.73 and a standard deviation of 0.21.

The original dataset additionally included multiple covariates, measuring the growing degree-days above zero, a moisture index, total solar radiation over the year, slope, topography, and elevation. In an attempt to identify the ecological gradient represented by the first latent variable, we *post-hoc* calculated correlation coefficients between the covariates and the first latent variable. From all covariates, elevation was most correlated with the first latent variable (a correlation coefficient of 0.93), though this was collinear with growing degree-days above zero and the moisture index. We additionally fitted two unconstrained linear GLLVMs with two latent variables, one of which included a random row intercept, and again calculated a correlation coefficient between the latent variables and elevation. The linear GLLVM without row intercept estimated the ecological gradient less successfully (highest correlation coefficient of −0.68), than when a row intercept was included (highest correlation coefficient of −0.92). To test more explicitly for the effect of elevation, we additionally fitted a quadratic GLLVM with elevation included as a covariate (both the linear and quadratic term), and with two latent variables. Including the covariate reduced the residual variance to 36% of that in the unconstrained model. The results presented here are from the unconstrained model, though the covariate effect of the second model is presented in appendix S5, Figure S3.

We examined groups of plants at the extremes of the gradient, i.e. plants that had optima of minus two or smaller, and plants with optima of two or larger, to further investigate whether the estimated latent variable from the quadratic GLLVM represented an elevation gradient. This approach allowed us to distinguish two groups of plants, the first indicative of lowlands (see Fig. 3). In contrast, plant species included on the opposite side of the latent variable were clearly indicative of alpine conditions. Here, we focus our inference on the alpine plants, as those are likely to be most affected by climate change (Walther *et al*. 2005). Of the alpine species, only two had confidence intervals for the quadratic coefficients that did not include zero: *Dryas octopetala* and *Sesleria caerulea. Dryas octopetala* had a maximum probability of occurrence of 0.35, a tolerance of 1.03, and an optimum at 2.05, and is as such predicted to be most prone to future changes in the environment. However, Figure 4 clearly shows some species that have more narrow tolerances, thus more specialised species are likely present in the dataset, though it was not possible to conclude this due to the confidence intervals of species quadratic coefficients crossing zero.

## Discussion

In this article, we extended the GLLVM approach of Hui *et al*. (2015), to estimate the niches of species with quadratic response curves, for unobserved ecological gradients. We fitted and performed inference for the quadratic GLLVM by extending the VA approach from Hui *et al*. (2017). The relation between latent variable models (i.e. unobserved ecological gradients) and ecological niches has been well described for classical ordination methods (ter Braak & Prentice 1988; Jongman *et al*. 1995), yet a method (either classical or model-based) to perform unconstrained (residual) ordination without limiting assumptions for species tolerances and maxima has not been available until now.

The similarity in responses of species to unobserved environments can be assed by examining tolerances, by examining an ordination diagrams for overlap in species distributions, or by using the residual correlation matrix. Determining if species exhibit fully quadratic curves in response to ecological gradients, whether tolerances are common for all species per ecological gradient, or if the equal tolerances assumption is suited for a dataset, comes down to a problem of model selection for the quadratic GLLVM. To that end, future research can further investigate approaches such as regularization (e.g., possibly extending the approach of Hui *et al*. 2018), hypothesis testing, or the use of confidence intervals of the quadratic coefficients. Similar to DCA, the quadratic GLLVM provides estimates of gradient length. But in contrast to DCA, where gradient length is a result of a heuristic rescaling of the ecological gradient, here it is calculated from the quadratic coefficients, which are estimated with approximate maximum likelihood.

For datasets with 50 species and 50 sites or more, the quadratic GLLVM accurately retrieved ecological gradients and species-specific parameters, though for continuous responses or counts it is possible to accurately estimate parameters with fewer species or sites. In general, when fitting the quadratic GLLVM to binary or ordinal responses, more information is required than for other data types (similarly as reported in Yee 2004). However, this is conditional on the information content in a dataset, and the number of required sites and species here should only be considered as a rough rule of thumb.

We studied the response of species to ecological gradients for hunting spiders in a Dutch dune ecosystem (van der Aart & Smeek-Enserink 1974), and for Swiss alpine plants (D’Amen *et al*. 2017), with use of the quadratic GLLVM. Various generalist species can be identified for both datasets, but as specialists are more likely to be affected by future changes in the environment, their identification is of critical importance to community ecology, to better focus recommendations for conservation efforts. We suggest that, for the hunting spiders dataset, *Arctosa perita*, and *Arctosa lutetiana* are most vulnerable to changes in the environment, and for the Swiss alpine plants dataset *Dryas octopetala* is most vulnerable to changes in the environment.

Modelling rare species is often difficult in community ecology as few ordination methods have the capability to explicitly do so. The quadratic GLLVM has great potential for community ecology, as it can simultaneously accommodate common (large tolerance and maxima i.e. a wide and high niche) and rare species (small tolerance and maxima i.e. a narrow and low niche) with the quadratic term. The quadratic GLLVM predicts species with unobserved optima, narrow niches, and small maxima will have the fewest observations. Since the quadratic GLLVM includes two species-specific parameters per latent variable, and thus requires more information in the data for accurate estimation of parameters than the linear GLLVM, it potentially requires a large dataset to include sufficient information on rare species. However, the example in this paper using the dataset of counts for hunting spiders (van der Aart & Smeek-Enserink 1974) shows that the quadratic GLLVM can be feasible to fit even to small datasets. An advantage of GLLVMs is their ability to use information from common species to improve estimation of parameters to describe the niches of rare species. However, without penalization or borrowing information for estimation from more abundant species, the parameters for species with few observations are not necessarily expected to be accurate.

The implementation of the quadratic GLLVM here is constrained to produce concave shapes only, though it could instead be used to estimate species minima rather than maxima. However, we did not do that here, as clear ecological foundations for such a model are lacking. An easy to use implementation based on the the `gllvm R` package (Niku *et al*. 2019b) is available on github (https://github.com/BertvanderVeen/gllvm-1/tree/goGLLVM), which will be included in the `gllvm R` package after publication.

## Acknowledgements

Manuela D’Amen kindly provided the elevation covariate for the Swiss Alpine plants dataset. B.V. was supported by a scholarship from the Research Council of Norway (grant numbver 272408/F40). F.K.C.H. was supported by two Australian Research Council Discovery grants.

## Authors contributions

B.V., K.A.H. and R.B.O. conceived the ideas. B.V., F.K.C.H. and R.B.O. designed the methodology. All authors contributed to the writing, reviewing and editing of the draft and gave final approval for publication.

## References

Björk, J.R., Hui, F.K.C., O’Hara, R.B. & Montoya, J.M. (2018). Uncovering the drivers of host-associated microbiota with joint species distribution modelling. Molecular Ecology, 27, 2714–2724.

Blanchet, F.G., Cazelles, K. & Gravel, D. (2020). Co-occurrence is not evidence of ecological interactions. Ecology Letters, 23, 1050–1063.

Borcard, D., Legendre, P. & Drapeau, P. (1992). Partialling out the Spatial Component of Ecological Variation. Ecology, 73, 1045–1055.

Burnham, K.P. & Anderson, D.R. (2002). Model Selection and Multimodel Inference: A Practical Information-Theoretic Approach, Secondn. Springer-Verlag, New York.

D’Amen, M., Mod, H.K., Gotelli, N.J. & Guisan, A. (2018). Disentangling biotic interactions, environmental filters, and dispersal limitation as drivers of species co-occurrence. Ecography, 41, 1233–1244.

D’Amen, M., Mod, H.K., Gotelli, N.J. & Guisan, A. (2017). Disentangling biotic interactions, environmental filters, and dispersal limitation as drivers of species co-occurrence. Dryad.

Damgaard, C., Hansen, R.R. & Hui, F.K.C. (2020). Model-based ordination of pin-point cover data: Effect of management on dry heathland. bioRxiv, 2020.03.05.980060.

Gabriel, K.R. (1971). The biplot graphic display of matrices with application to principal component analysis. Biometrika, 58, 453–467.

Gauch, H.G. (1982). Multivariate Analysis in Community Ecology. Cambridge University Press, Cambridge.

Halvorsen, R. (2012). A gradient analytic perspective on distribution modelling. Sommerfeltia, 35, 1–165.

Hill, M.O. & Gauch, H.G. (1980). Detrended Correspondence Analysis: An Improved Ordination Technique. Classification and Ordination: Symposium on advances in vegetation science, Nijmegen, The Netherlands, May 1979 (ed E. van der Maarel), pp. 47–58. Advances in vegetation science. Springer Netherlands, Dordrecht.

Hui, F.K.C. (2016). Boral Bayesian Ordination and Regression Analysis of Multivariate Abundance Data in r. Methods in Ecology and Evolution, 7, 744–750.

Hui, F.K.C., Tanaka, E. & Warton, D.I. (2018). Order selection and sparsity in latent variable models via the ordered factor LASSO. Biometrics, 74, 1311–1319.

Hui, F.K.C., Taskinen, S., Pledger, S., Foster, S.D. & Warton, D.I. (2015). Model-based approaches to unconstrained ordination. Methods in Ecology and Evolution, 6, 399–411.

Hui, F.K.C., Warton, D.I., Ormerod, J.T., Haapaniemi, V. & Taskinen, S. (2017). Variational Approximations for Generalized Linear Latent Variable Models. Journal of Computational and Graphical Statistics, 26, 35–43.

Inoue, K., Stoeckl, K. & Geist, J. (2017). Joint species models reveal the effects of environment on community assemblage of freshwater mussels and fishes in European rivers. Diversity and Distributions, 23, 284–296.

Jamil, T. & ter Braak, C.J.F. (2013). Generalized linear mixed models can detect unimodal species-environment relationships. PeerJ, 1, e95.

Jongman, R., ter Braak, C. & van Tongeren, O. (Eds.). (1995). Data analysis in community and landscape ecology. Cambridge university press, Cambridge.

Khatri, C.G. (1980). 14 Quadratic forms in normal variables. Handbook of Statistics, pp. 443–469.Analysis of Variance. Elsevier.

Kristensen, K., Nielsen, A., Berg, C.W., Skaug, H. & Bell, B. (2016). TMB: Automatic Differentiation and Laplace Approximation. Journal of Statistical Software, 70. Retrieved from http://arxiv.org/abs/1509.00660

Lacoste, É., Weise, A.M., Lavoie, M.-F., Archambault, P. & McKindsey, C.W. (2019). Changes in infaunal assemblage structure influence nutrient fluxes in sediment enriched by mussel biodeposition. Science of The Total Environment, 692, 39–48.

MacArthur, R. & Levins, R. (1967). The limiting similarity, convergence, and divergence of coexisting species. The American Naturalist, 101, 377–385.

Nakagawa, S. & Schielzeth, H. (2013). A general and simple method for obtaining R2 from generalized linear mixed-effects models. Methods in Ecology and Evolution, 4, 133–142.

Niku, J., Brooks, W., Herliansyah, R., Hui, F.K.C., Taskinen, S. & Warton, D.I. (2019a). Efficient estimation of generalized linear latent variable models. PLOS ONE, 14, e0216129.

Niku, J., Brooks, W., Herliansyah, R., Hui, F.K.C., Taskinen, S. & Warton, D.I. (2020). Gllvm: Generalized linear latent variable models.

Niku, J., Hui, F.K.C., Taskinen, S. & Warton, D.I. (2019b). Gllvm: Fast analysis of multivariate abundance data with generalized linear latent variable models in r. Methods in Ecology and Evolution, 10, 2173–2182.

Oksanen, J. & Tonteri, T. (1995). Rate of compositional turnover along gradients and total gradient length. Journal of Vegetation Science, 6, 815–824.

Ovaskainen, O., Tikhonov, G., Norberg, A., Blanchet, F.G., Duan, L., Dunson, D., Roslin, T. & Abrego, N. (2017). How to make more out of community data? A conceptual framework and its implementation as models and software. Ecology Letters, 20, 561–576.

Peres-Neto, P.R. & Jackson, D.A. (2001). How well do multivariate data sets match? The advantages of a Procrustean superimposition approach over the Mantel test. Oecologia, 129, 169–178.

Pollock, L.J., Tingley, R., Morris, W.K., Golding, N., O’Hara, R.B., Parris, K.M., Vesk, P.A. & McCarthy, M.A. (2014). Understanding co-occurrence by modelling species simultaneously with a Joint Species Distribution Model (JSDM). Methods in Ecology and Evolution, 5, 397–406.

ter Braak, C.J. (1986). Canonical Correspondence Analysis: A New Eigenvector Technique for Multivariate Direct Gradient Analysis. Ecology, 67, 1167–1179.

ter Braak, C.J.F. (1985). Correspondence Analysis of Incidence and Abundance Data: Properties in Terms of a Unimodal Response Model. Biometrics, 41, 859–873.

ter Braak, C.J.F. & Prentice, I.C. (1988). A Theory of Gradient Analysis. Advances in Ecological Research (eds M. Begon, A.H. Fitter, E.D. Ford & A. Macfadyen), pp. 271–317. Academic Press.

Tikhonov, G., Ovaskainen, O., Oksanen, J., de Jonge, M., Opedal, O. & Dallas, T. (2020). Hmsc: Hierarchical model of species communities.

Tobler, M.W., Kéry, M., Hui, F.K.C., Guillera-Arroita, G., Knaus, P. & Sattler, T. (2019). Joint species distribution models with species correlations and imperfect detection. Ecology, 100, e02754.

van der Aart, P. & Smeek-Enserink, N. (1974). Correlations between distributions of hunting spiders (Lycosidae, Ctenidae) and environmental characteristics in a dune area. Netherlands Journal of Zoology, 25, 1–45.

Walker, S.C. & Jackson, D.A. (2011). Random-effects ordination: Describing and predicting multivariate correlations and co-occurrences. Ecological Monographs, 81, 635–663.

Walther, G.-R., Beißner, S. & Burga, C.A. (2005). Trends in the upward shift of alpine plants. Journal of Vegetation Science, 16, 541–548.

Wang, Y., Naumann, U., Wright, S.T. & Warton, D.I. (2012). Mvabund an R package for model-based analysis of multivariate abundance data. Methods in Ecology and Evolution, 3, 471–474.

Warton, D.I., Blanchet, F.G., O’Hara, R.B., Ovaskainen, O., Taskinen, S., Walker, S.C. & Hui, F.K.C. (2015). So Many Variables: Joint Modeling in Community Ecology. Trends in Ecology & Evolution, 30, 766–779.

Wehrden, H.V., Hanspach, J., Bruelheide, H. & Wesche, K. (2009). Pluralism and diversity: Trends in the use and application of ordination methods 1990-2007. Journal of Vegetation Science, 20, 695–705.

Yee, T.W. (2004). A New Technique for Maximum-Likelihood Canonical Gaussian Ordination. Ecological Monographs, 74, 685–701.

Zurell, D., Pollock, L.J. & Thuiller, W. (2018). Do joint species distribution models reliably detect interspecific interactions from co-occurrence data in homogenous environments? Ecography, 41, 1812–1819.

Zurell, D., Zimmermann, N.E., Gross, H., Baltensweiler, A., Sattler, T. & Wüest, R.O. (2020). Testing species assemblage predictions from stacked and joint species distribution models. Journal of Biogeography, 47, 101–113.

Økland, R.H. (1999). On the variation explained by ordination and constrained ordination axes. Journal of Vegetation Science, 10, 131–136.

Økland, R.H. & Eilertsen, O. (1994). Canonical Correspondence Analysis with variation partitioning: Some comments and an application. Journal of Vegetation Science, 5, 117–126.

