## Supplementary information for "Model-based ordination for species with unequal niche widths"

#### Appendix S1: Residual covariance

Here, we provide a derivation for the residual covariance of a GLLVM with quadratic latent variables. For a vector of latent variables $\boldsymbol{z}_i$, which is assumed to follow a multivariate standard normal distribution i.e. $\boldsymbol{z}_i \sim \mathcal{N}(\boldsymbol{0}, \boldsymbol{I})$, where sites are assumed to be independent, and for the linear predictor $\eta_{ij} = C_{ij} + \boldsymbol{z}_i^\top \boldsymbol{\gamma}_j - \boldsymbol{z}_i^\top \boldsymbol{D}_j \boldsymbol{z}_i$, where $C_{ij}$ is a general quantity that is constant with respect to the latent variables, the entries of the residual covariance matrix $\boldsymbol{\Sigma}$ for species $j, l = 1 \dots p$, are given by:

$$
\begin{aligned}
\text{cov}(\boldsymbol{z}_i^\top \boldsymbol{\gamma}_j - \boldsymbol{z}_i^\top \boldsymbol{D}_j \boldsymbol{z}_i, \boldsymbol{z}_i^\top \boldsymbol{\gamma}_l - \boldsymbol{z}_i^\top \boldsymbol{D}_l \boldsymbol{z}_i) =& \text{cov}(\boldsymbol{z}_i^\top \boldsymbol{\gamma}_j, \boldsymbol{z}_i^\top \boldsymbol{\gamma}_l) \\
&+ \text{cov}(\boldsymbol{z}_i^\top \boldsymbol{\gamma}_j, -\boldsymbol{z}_i^\top \boldsymbol{D}_l \boldsymbol{z}_i) \\
&+ \text{cov}(\boldsymbol{z}_i^\top \boldsymbol{\gamma}_l, -\boldsymbol{z}_i^\top \boldsymbol{D}_j \boldsymbol{z}_i) \\
&+ \text{cov}(-\boldsymbol{z}_i^\top \boldsymbol{D}_j \boldsymbol{z}_i, -\boldsymbol{z}_i^\top \boldsymbol{D}_l \boldsymbol{z}_i).
\end{aligned}
$$

Since the third order central moments of the multivariate normal distribution are zero, then we only have to calculate the first and last term,

$$\begin{aligned}\operatorname{cov}(\boldsymbol{z}_i^\top \boldsymbol{\gamma}_j - \boldsymbol{z}_i^\top \boldsymbol{D}_j \boldsymbol{z}_i, \boldsymbol{z}_i^\top \boldsymbol{\gamma}_l - \boldsymbol{z}_i^\top \boldsymbol{D}_l \boldsymbol{z}_i) =& \operatorname{cov}(\boldsymbol{z}_i^\top \boldsymbol{\gamma}_j, \boldsymbol{z}_i^\top \boldsymbol{\gamma}_l) \\ &+ \operatorname{cov}(-\boldsymbol{z}_i^\top \boldsymbol{D}_j \boldsymbol{z}_i, -\boldsymbol{z}_i^\top \boldsymbol{D}_l \boldsymbol{z}_i) \\ =& \boldsymbol{\gamma}_j^\top \boldsymbol{\gamma}_l + 2\operatorname{tr}(\boldsymbol{D}_j \boldsymbol{D}_l). \end{aligned} \tag{1}$$

### Appendix S2: Variational approximations

In the derivation of the VA log-likelihoods below, we let $C_{ij}$ generically denote a quantity that is constant with respect to the latent variables $\boldsymbol{z}_i$. For example, species-specific intercepts, for $i = 1 \dots n$ sites and $j = 1 \dots p$ species. We start by defining the linear predictor:

$$\eta_{ij} = C_{ij} + \boldsymbol{z}_i^\top \boldsymbol{\gamma}_j - \boldsymbol{z}_i^\top \boldsymbol{D}_j \boldsymbol{z}_i,$$

where $\boldsymbol{\gamma}_j$ is a vector of species coefficients for the linear term of the $q = 1 \dots d$ latent variables, $\boldsymbol{D}_j$ is a species-specific positive-definite diagonal matrix of size $d \times d$. Below, $\boldsymbol{\Theta}$ is a vector including all model parameters i.e. $\boldsymbol{\Theta} = (C_{11} \dots C_{ij}, \gamma_{11} \dots \gamma_{jq}, D_{111} \dots D_{jqq})^\top$, including any nuisance parameters where applicable, such as cutoffs $\boldsymbol{\zeta}_{jk}$ for ordinal responses, and dispersion parameters $\phi_j$ for the Gaussian, gamma, and negative-binomial responses.

#### Variational approximations

The GLLVM log-likelihood is:

$$\mathcal{L}(\boldsymbol{\Theta}) = \sum_{i=1}^{n} \log \bigg\{ \int \prod_{j=1}^{p} f\Big(y_{ij}|\boldsymbol{z}_i, \boldsymbol{\Theta}\Big) h\Big(\boldsymbol{z}_i\Big) d\boldsymbol{z}_i \bigg\}, \tag{2}$$

where $\boldsymbol{z}_i \sim \mathcal{N}(\boldsymbol{0}, \mathbf{I})$, and where $f(y_{ij}|\boldsymbol{z}_i, \boldsymbol{\Theta})$ is a GLM-type distribution (e.g. in the exponential family). Variational approximations (VA) applies Jensen's inequality to construct a lower bound for the marginal log-likelihood (see; Hui et al., 2017) giving:

$$\log \mathcal{L}_{VA}(\boldsymbol{\Theta}, \boldsymbol{\xi}) = \sum_{i=1}^{n} \sum_{j=1}^{p} \bigg\{ \int \log\bigg( \frac{f(y_{ij}|\boldsymbol{z}_i, \boldsymbol{\Theta}) h(\boldsymbol{z}_i)}{\mathrm{q}(\boldsymbol{z}_i|\boldsymbol{a}_i, \boldsymbol{A}_i)} \bigg) \mathrm{q}(\boldsymbol{z}_i|\boldsymbol{a}_i, \boldsymbol{A}_i) d\boldsymbol{z}_i \bigg\}, \tag{3}$$

where $\mathrm{q}(\boldsymbol{z}_i|\boldsymbol{a}_i, \boldsymbol{A}_i)$ is a variational distribution of the latent variables, which we assume to be multivariate normal with mean $\boldsymbol{a}_i$ and covariance matrix $\boldsymbol{A}_i$, so that $\boldsymbol{\xi} = \{a_{11} \dots a_{iq}, \mathrm{vech}(\boldsymbol{A}_i)\}^\top$. The variational distribution serves as a closed form approximation to the true posterior distribution of the latent variables. The variational distribution of the latent variables can be found by following equation (5) from Ormerod and Wand (2010). However, a simpler parametric form for the variational distribution can be assumed to ensure a tractable likelihood. The Kullback-Leibler divergence is minimized, so that the combination of a multivariate normal distribution with the variational parameters best represent the true posterior distribution. The VA log-likelihood is then calculated by taking expectations over the components of equation (3):

$$\log \mathcal{L}_{VA}(\boldsymbol{\Theta}, \boldsymbol{\xi}) = \sum_{i=1}^{n}\sum_{j=1}^{p} \text{E}[\log\{f(y_{ij}|\boldsymbol{z}_i, \boldsymbol{\Theta})\}\text{q}\{\boldsymbol{z}_i|\boldsymbol{a}_i, \boldsymbol{A}_i\}] - \sum_{i=1}^{n} \text{E}\{\log \text{q}(\boldsymbol{z}_i|\boldsymbol{a}_i, \boldsymbol{A}_i)\}, \tag{4}$$

Expectations for the second term in equation (4) follow the standard result:

$$\text{E}[\log\{\text{q}(\boldsymbol{z}_i|\boldsymbol{a}_i, \boldsymbol{A}_i)\}] = \frac{1}{2}\log\det(\boldsymbol{A}_i) - \text{tr}(\boldsymbol{A}_i) - \boldsymbol{a}_i^\top \boldsymbol{a}_i.$$

Additionally, taking expectations over the linear predictor from above, with respect to the variational distribution of the latent variables, gives:

$$\tilde{\boldsymbol{\eta}}_{ij} = C_{ij} + \text{E}(\boldsymbol{z}_i^\top)\boldsymbol{\gamma}_j + \text{E}(\boldsymbol{z}_i^\top \boldsymbol{D}_j \boldsymbol{z}_i) = C_{ij} + \boldsymbol{a}_i^\top \boldsymbol{\gamma_j} - \boldsymbol{a}_i^\top \boldsymbol{D}_j \boldsymbol{a}_i - \text{tr}(\boldsymbol{D}_j \boldsymbol{A}_i).$$

These two results above are valid for all distributions, so that the general VA log-likelihood for the quadratic GLLVM is:

$$\log \mathcal{L}_{VA}(\boldsymbol{\Theta}, \boldsymbol{\xi}) = \sum_{i=1}^{n}\sum_{j=1}^{p}\left[\frac{y_{ij}\tilde{\eta}_{ij} - \text{E}\{\text{b}(\eta_{ij})\}}{\text{a}\{\phi_j\}} + c\{y_{ij}, \phi_j\}\right] + \frac{1}{2}\sum_{i=1}^{n}\left\{\log\det\left(\boldsymbol{A}_i\right) - \text{tr}\left(\boldsymbol{A}_i\right) - \boldsymbol{a}_i^\top \boldsymbol{a}_i\right\}, \tag{5}$$

where $a(\cdot)$, $b(\cdot)$, and $c(\cdot)$ are known functions. Calculating $\int \text{b}(\eta_{ij})h(\boldsymbol{z}_i)d\boldsymbol{z}_i$, is most challenging as the solution has to be derived separately for most distributions. Below follow the derivations for the Gaussian, Poisson, negative-binomial, Bernoulli, ordinal and gamma likelihoods.

#### Gaussian: continuous responses

The joint log-likelihood for Gaussian distributed responses is:

$$\mathcal{L}(\boldsymbol{\Theta}) = \sum_{i=1}^{n}\sum_{j=1}^{p}\left\{-\frac{1}{2}\log\left(\sigma_j^2\right) - \frac{1}{2\sigma_j^2}\left(y_{ij} - \eta_{ij}\right)^2\right\} - \frac{1}{2}\sum_{i=1}^{n}\boldsymbol{z}_i^\top \boldsymbol{z}_i, \tag{6}$$

where terms constant as a function of the parameters have been omitted. The VA log-likelihood is derived by taking expectations over the components of equation (3) with respect to the latent variables:

$$\text{E}[\log\{f(y_{ij}|\boldsymbol{z}_i, \boldsymbol{\Theta})h(\boldsymbol{z}_i)\}] = \sum_{i=1}^{n}\sum_{j=1}^{p}\left[-\frac{1}{2}\log\left\{\sigma_j^2\right\} - \frac{1}{2\sigma_j^2}\text{E}\left\{\left(y_{ij} - \eta_{ij}\right)^2\right\}\right] - \frac{1}{2}\sum_{i=1}^{n}\left\{\text{tr}\left(\boldsymbol{A}_i\right) + \boldsymbol{a}_i^\top \boldsymbol{a}_i\right\}$$

$$\text{E}[\log\{\text{q}(\boldsymbol{z}_i|\boldsymbol{a}_i, \boldsymbol{A}_i)\}] = -\frac{1}{2}\sum_{i=1}^{n}\log\det(2e\pi\boldsymbol{A}_i),$$

where terms constant with respect to the parameters have been omitted. The term $\mathrm{E}\{(y_{ij} - \eta_{ij})^2\}$ can instead be written and expanded as:

$$\mathrm{E}\{(y_{ij} - \eta_{ij})^2\} = \mathrm{E}\{(y_{ij} - \tilde{\eta}_{ij} + \tilde{\eta}_{ij} - \eta_{ij})^2\}$$

$$\mathrm{E}\{(y_{ij} - \tilde{\eta}_{ij})^2\} + 2\mathrm{E}\{(y_{ij} - \tilde{\eta}_{ij})(\tilde{\eta}_{ij} - \eta_{ij})\} + \mathrm{E}\{(\tilde{\eta}_{ij} - \eta_{ij})^2\}.$$

As the first term does not require taking expectations, and since the second term is zero, only the third term has to be calculated, so that:

$$\mathrm{E}\{(\tilde{\eta}_{ij} - \eta_{ij})^2\} = \mathrm{var}(\eta_{ij})$$

$$\begin{aligned}
\mathrm{var}(\eta_{ij}) &= \mathrm{var}(C_{ij} + \boldsymbol{z}_i^\top \boldsymbol{\gamma}_j - \boldsymbol{z}_i^\top \boldsymbol{D}_j \boldsymbol{z}_i) \\
&= \mathrm{var}(\boldsymbol{z}_i^\top \boldsymbol{\gamma}_j) + \mathrm{var}(\boldsymbol{z}_i^\top \boldsymbol{D}_j \boldsymbol{z}_i) - 2\mathrm{cov}(\boldsymbol{z}_i^\top \boldsymbol{\gamma}_j, \boldsymbol{z}_i^\top \boldsymbol{D}_j \boldsymbol{z}_i) \\
&= \mathrm{tr}(\boldsymbol{\gamma}_j \boldsymbol{\gamma}_j^\top \boldsymbol{A}_i) + 2\mathrm{tr}(\boldsymbol{D}_j \boldsymbol{A}_i \boldsymbol{D}_j \boldsymbol{A}_i) + 4\boldsymbol{a}_i^\top \boldsymbol{D}_j \boldsymbol{A}_i \boldsymbol{D}_j \boldsymbol{a}_i - 2\mathrm{cov}(\boldsymbol{z}_i^\top \boldsymbol{\gamma}_j, \boldsymbol{z}_i^\top \boldsymbol{D}_j \boldsymbol{z}_i).
\end{aligned}$$

For the last term:

$$\begin{aligned}
\mathrm{cov}(\boldsymbol{z}_i^\top \boldsymbol{\gamma}_j, \boldsymbol{z}_i^\top \boldsymbol{D}_j \boldsymbol{z}_i) &= \mathrm{cov}\{\boldsymbol{z}_i^\top \boldsymbol{\gamma}_j - \mathrm{E}(\boldsymbol{z}_i^\top \boldsymbol{\gamma}_j), \boldsymbol{z}_i^\top \boldsymbol{D}_j \boldsymbol{z}_i - \mathrm{E}(\boldsymbol{z}_i^\top \boldsymbol{D}_j \boldsymbol{z}_i)\} \\
&= \mathrm{cov}\{\sum_{k=1}^{d} \gamma_{jk}(u_{ik} - a_{ik}), \sum_{l=1}^{d} D_{jll} u_{il}^2 - D_{jll}\mathrm{E}(u_{il}^2)\} \\
&= \sum_{k,l=1}^{d} \mathrm{cov}\{\gamma_{jk}(u_{ik} - a_{ik}), D_{jll} u_{il}^2 - D_{jll}\mathrm{E}(u_{il}^2)\} \\
&= \sum_{k,l=1}^{d} \mathrm{E}[\gamma_{jk}\{u_{ik} - a_{ik}\}\{D_{jll} u_{il}^2 - D_{jll}\mathrm{E}(u_{il}^2)\}] \\
&= \sum_{k,l=1}^{d} \mathrm{E}\{\gamma_{jk}(u_{ik} - a_{ik}) D_{jll} u_{il}^2\} \\
&= \sum_{k,l=1}^{d} \gamma_{jk} D_{jll} \mathrm{E}\{(u_{ik} - a_{ik})(u_{il} - a_{il} + a_{il})^2\} \\
&= \sum_{k,l=1}^{d} \gamma_{jk} D_{jll} \mathrm{E}\{(u_{ik} - a_{ik})(u_{il} - a_{il})^2 + 2a_{il}(u_{ik} - a_{ik})(u_{il} - a_{il})\} \\
&= \sum_{k,l=1}^{d} 2\gamma_{jk} D_{jll} a_{il} \mathrm{E}\{(u_{ik} - a_{ik})(u_{il} - a_{il})\} \\
&= 2\sum_{k,l=1}^{d} \gamma_{jk} D_{jll} a_{il} A_{ikl} \\
&= 2\boldsymbol{a}_i^\top \boldsymbol{D}_j \boldsymbol{A}_i \boldsymbol{\gamma}_j.
\end{aligned} \tag{9}$$

Thus, the Gaussian VA log-likelihood for the quadratic model is:

$$\mathcal{L}_{VA}(\boldsymbol{\Theta}, \boldsymbol{\xi}) = \sum_{i=1}^{n}\sum_{j=1}^{p}\Bigg[-\frac{1}{2}\log\Big\{\sigma_j^2\Big\} - \frac{1}{2\sigma_j^2}\Big\{y_{ij}^2 + \tilde{\eta}_{ij}^2 - 2y_{ij}\tilde{\eta}_{ij} + \mathrm{tr}\Big(\boldsymbol{\gamma}_j\boldsymbol{\gamma}_j^\top\boldsymbol{A}_i\Big) + 2\mathrm{tr}\Big(\boldsymbol{D}_j\boldsymbol{A}_i\boldsymbol{D}_j\boldsymbol{A}_i\Big) + 4\boldsymbol{a}_i^\top\boldsymbol{D}_j\boldsymbol{A}_i\boldsymbol{D}_j\boldsymbol{a}_i - 4\boldsymbol{a}_i^\top\boldsymbol{D}_j\boldsymbol{A}_i\boldsymbol{\gamma}_j\Big\}\Bigg] + \frac{1}{2}\sum_{i=1}^{n}\Big\{\log\det\Big(\boldsymbol{A}_i\Big) - \mathrm{tr}\Big(\boldsymbol{A}_i\Big) - \boldsymbol{a}_i^\top\boldsymbol{a}_i\Big\} \quad (10)$$

### Bernoulli: presence-absence responses

We model presence-absence data with a probit-link function and a Bernoulli distribution, giving the joint log-likelihood:

$$\mathcal{L}(\boldsymbol{\Theta}) = \sum_{i=1}^{n}\sum_{j=1}^{p}\Bigg[y_{ij}\log\Big\{\mathrm{I}\Big(\nu_{ij} \geq 0\Big)\Big\} + \Big\{1 - y_{ij}\Big\}\log\Big\{\mathrm{I}\Big(\nu_{ij} < 0\Big)\Big\} - \frac{1}{2}\Big\{\nu_{ij} - \eta_{ij}\Big\}^2\Bigg] - \frac{1}{2}\sum_{i=1}^{n}\boldsymbol{z}_i^\top\boldsymbol{z}_i, \quad (11)$$

where terms constant as a function of the parameters have been omitted. Here, $v_{ij} \sim \mathcal{N}(\eta_{ij}, 1)$ is an auxiliary variable to aid integration, with indicator function $\mathrm{I}(\cdot)$. As in Hui et al. (2017), and by equation (5) in Ormerod and Wand (2010), the optimal variational distribution for the auxiliary variable is calculated by taking expectations over equation (11) with respect to $\nu_{ij}$:

$$\begin{aligned} \log\{\mathrm{q}(\nu_{ij})\} &\propto \mathrm{E}\{\mathcal{L}(\boldsymbol{\Theta})\} \\ \log\{\mathrm{q}(\nu_{ij})\} &\propto \log\{\mathrm{I}(\nu_{ij} > 0)\} - \frac{1}{2}\{\nu_{ij} - \mathrm{E}(\eta_{ij})\}^2, \qquad y_{ij} = 1 \\ \log\{\mathrm{q}(\nu_{ij})\} &\propto \log\{\mathrm{I}(\nu_{ij} < 0)\} - \frac{1}{2}\{\nu_{ij} - \mathrm{E}(\eta_{ij})\}^2, \qquad y_{ij} = 0. \end{aligned} \quad (12)$$

Terms constant as a function of the parameters have been omitted. The second term of both lines in equation (12) expands to $\nu_{ij}\mathrm{E}(\eta_{ij}) - \frac{1}{2}\nu_{ij}^2 - \frac{1}{2}\mathrm{E}(\eta_{ij}^2)$, showing that the distribution of $\nu_{ij}$ does not depend on $\eta_{ij}^2$. As equation (12) is still quadratic after taking expectations, the optimal distribution for the auxiliary variable is truncated normal with location parameter $\mathrm{E}(\eta_{ij}) = \tilde{\eta}_{ij}$ and scale parameter one, as in Hui et al. (2017). The distribution has limits $(0,\infty)$ and $(-\infty,0)$ for $y_{ij} = 1$ and $y_{ij} = 0$ respectively.

Due to the inclusion of the auxiliary variable, the components of the VA log-likelihood are now:

$$\log\mathcal{L}_{VA} = \sum_{i=1}^{n}\sum_{j=1}^{p}\mathrm{E}[\log\{f(y_{ij}|\boldsymbol{z}_i, \boldsymbol{\Theta})h(\boldsymbol{z}_i)\}] - \mathrm{E}\{\log\mathrm{q}(\nu_{ij})\} - \sum_{i=1}^{n}\mathrm{E}\{\log\mathrm{q}(\boldsymbol{z}_i|\boldsymbol{a}_i, \boldsymbol{A}_i)\}. \quad (13)$$

Then, taking expectations over each of the components in equation (13),

$$\mathrm{E}[\log\{f(y_{ij}|\boldsymbol{z}_i,\boldsymbol{\Theta})h(\boldsymbol{z}_i)\}] = \sum_{i=1}^{n}\sum_{j=1}^{p}\mathrm{E}\Big[y_{ij}\log\Big\{\mathrm{I}\Big(\nu_{ij}>0\Big)\Big\}+\Big\{1-y_{ij}\Big\}\log\Big\{\mathrm{I}\Big(\nu_{ij}<0\Big)\Big\}\Big]-\frac{1}{2}\mathrm{E}\Big[\Big\{\nu_{ij}-\eta_{ij}\Big\}^2\Big] \\ -\frac{1}{2}\sum_{i=1}^{n}\Big\{\mathrm{tr}\Big(\boldsymbol{A}_i\Big)+\boldsymbol{a}_i^\top\boldsymbol{a}_i\Big\} \tag{14}$$

$$\mathrm{E}[\log\{\mathrm{q}(\nu_{ij})\}] = \mathrm{E}\Big[y_{ij}\log\Big\{\mathrm{I}\Big(\nu_{ij}>0\Big)\Big\}+\Big(1-y_{ij}\Big)\log\Big\{\mathrm{I}\Big(\nu_{ij}<0\Big)\Big\}\Big]-\frac{1}{2}\mathrm{E}\Big[\Big\{\nu_{ij}-\tilde{\eta}_{ij}\Big\}^2\Big] \\ -y_{ij}\log\Big\{\Phi\Big(\tilde{\eta}_{ij}\Big)\Big\}-\Big(1-y_{ij}\Big)\log\Big\{1-\Phi\Big(\tilde{\eta}_{ij}\Big)\Big\}$$

$$\mathrm{E}[\log\{\mathrm{q}(\boldsymbol{z}_i|\boldsymbol{a}_i,\boldsymbol{A}_i)\}] = -\frac{1}{2}\sum_{i=1}^{n}\log\det(2e\pi\boldsymbol{A}_i),$$

where terms constant as a function of the parameters have been omitted. The term $\mathrm{E}[\{\nu_{ij}-\eta_{ij}\}^2]$ can be rewritten to simplify taking expectations, in the following way:

$$\mathrm{E}\{(\nu_{ij}-\eta_{ij})^2\} = \mathrm{E}\{(\nu_{ij}-\tilde{\eta}_{ij}+\tilde{\eta}_{ij}-\eta_{ij})^2\}$$

$$\mathrm{E}\{(\nu_{ij}-\tilde{\eta}_{ij})^2\}+2\mathrm{E}\{(\nu_{ij}-\tilde{\eta}_{ij})(\tilde{\eta}_{ij}-\eta_{ij})\}+\mathrm{E}\{(\tilde{\eta}_{ij}-\eta_{ij})^2\}.$$

The first term cancels with the same in the first line of equation (14), the second term is zero under the assumption of independent variational distributions and because $\eta_{ij}$ is not a function of $\nu_{ij}$. The solution to the last term is given in equation (9).

Thus, the Bernoulli VA log-likelihood for the quadratic model is:

$$\mathcal{L}_{VA}(\boldsymbol{\Theta},\boldsymbol{\xi}) = \sum_{i=1}^{n}\sum_{j=1}^{p}\Big[y_{ij}\log\Big\{\Phi\Big(\tilde{\eta}_{ij}\Big)\Big\}+\Big\{1-y_{ij}\Big\}\log\Big\{1-\Phi\Big(\tilde{\eta}_{ij}\Big)\Big\}-\frac{1}{2}\mathrm{tr}\Big\{\boldsymbol{\gamma}_j\boldsymbol{\gamma}_j^\top\boldsymbol{A}_i\Big\}-\mathrm{tr}\Big\{\boldsymbol{D}_j\boldsymbol{A}_i\boldsymbol{D}_j\boldsymbol{A}_i\Big\} \\ -2\boldsymbol{a}_i^\top\boldsymbol{D}_j\boldsymbol{A}_i\boldsymbol{D}_j\boldsymbol{a}_i+2\boldsymbol{a}_i^\top\boldsymbol{D}_j\boldsymbol{A}_i\boldsymbol{\gamma}_j\Big]+\frac{1}{2}\sum_{i=1}^{n}\Big\{\log\det\Big(\boldsymbol{A}_i\Big)-\mathrm{tr}\Big(\boldsymbol{A}_i\Big)-\boldsymbol{a}_i^\top\boldsymbol{a}_i\Big\}. \tag{16}$$

#### Ordinal: ordered responses

The ordered response model follows from an extension of the Bernoulli VA log-likelihood for multiple categories. We define a matrix of cutoffs $\boldsymbol{\zeta}_{jk}$, which serves to introduce order to the probability of occurrence in each of the $k = 1 \dots K$ categories.

The first cutoff is set to zero for reasons of parameter identifiability, and so that $\zeta_{j0} = -\infty$ and $\zeta_{jK} = \infty$, ensuring that the probability of occurrence in the first category is at most $\frac{1}{2}$, corresponding with the Bernoulli distribution when $K = 2$.

With the assumption of species-specific cutoffs as above, comes the requirement of more than one observation per species and category. This requirement can be relaxed, by noting that the categories per species are arbitrary (as they are indexes only), so that they may be renumbered to exclude categories that lack any observations (i.e. missing categories). To improve the useability of the ordinal model, we here introduce an additional, simpler, parameterization where we allow $\zeta_{jk} = \zeta_k$, i.e. species-common cutoffs, so that $\zeta_0 = -\infty$ and $\zeta_K = \infty$. This relaxes the requirement of at least one observation per category per species to at least one observation per category in the whole dataset, which we deem more realistic for real world data.

In either case, the variational distribution for auxiliary variable $\nu_{ij}$ is truncated normal, but with limits $\zeta_{k-1}, \zeta_k$ for $y_{ijk} > 0$ in the species-common cutoff case. Additionally, we define $y_{ijk}$ as a K-dimensional array, so that the ordinal VA log-likelihood for the quadratic model is:

$$\begin{aligned}\mathcal{L}_{VA}(\boldsymbol{\Theta}, \boldsymbol{\xi}) = \sum_{i=1}^{n}\sum_{j=1}^{p}\sum_{k=1}^{K_j}\Bigg[y_{ijk}\log\Bigg\{\Phi\Big(\zeta_{jk} - \tilde{\eta}_{ij}\Big) - \Phi\Big(\zeta_{jk-1} - \tilde{\eta}_{ij}\Big)\Bigg\} - \frac{1}{2}\mathrm{tr}\Big\{\boldsymbol{\gamma}_j\boldsymbol{\gamma}_j^\top\boldsymbol{A}_i\Big\} - \mathrm{tr}\Big\{\boldsymbol{D}_j\boldsymbol{A}_i\boldsymbol{D}_j\boldsymbol{A}_i\Big\} \\ - 2\boldsymbol{a}_i^\top\boldsymbol{D}_j\boldsymbol{A}_i\boldsymbol{D}_j\boldsymbol{a}_i + 2\boldsymbol{a}_i^\top\boldsymbol{D}_j\boldsymbol{A}_i\boldsymbol{\gamma}_j\Bigg] + \frac{1}{2}\sum_{i=1}^{n}\Bigg\{\log\det\Big(\boldsymbol{A}_i\Big) - \mathrm{tr}\Big(\boldsymbol{A}_i\Big) - \boldsymbol{a}_i^\top\boldsymbol{a}_i\Bigg\}.\end{aligned} \tag{17}$$

#### Poisson: counted responses

The joint log-likelihood for Poisson distributed responses with a log-link function is:

$$\mathcal{L}(\boldsymbol{\Theta}) = \sum_{i=1}^{n}\sum_{j=1}^{p}\Big\{y_{ij}\eta_{ij} - \exp\Big(\eta_{ij}\Big)\Big\} - \frac{1}{2}\sum_{i=1}^{n}\boldsymbol{z}_i^\top\boldsymbol{z}_i, \tag{18}$$

where terms constant as a function of the parameters have been omitted. The VA log-likelihood is derived by taking expectations over the components of equation (4) with respect to the latent variables:

$$\mathrm{E}[\log\{f(y_{ij}|\boldsymbol{z}_i, \boldsymbol{\Theta})h(\boldsymbol{z}_i)\}] = \sum_{i=1}^{n}\sum_{j=1}^{p}\Bigg[y_{ij}\mathrm{E}\Big\{\eta_{ij}\Big\} - \mathrm{E}\Big\{\exp\Big(\eta_{ij}\Big)\Big\}\Bigg] - \frac{1}{2}\sum_{i=1}^{n}\Big\{\mathrm{tr}(\boldsymbol{A}_i) + \boldsymbol{a}_i^\top\boldsymbol{a}_i\Big\} \tag{19a}$$

$$\mathrm{E}[\log\{\mathrm{q}(\boldsymbol{z}_i|\boldsymbol{a}_i, \boldsymbol{A}_i)\}] = -\frac{1}{2}\sum_{i=1}^{n}\log\det(2e\pi\boldsymbol{A}_i),$$

where terms constant as a function of the parameters have been omitted. To retrieve a closed form for the integration of $\mathrm{E}\{\exp(\eta_{ij})\}$, we rewrite the quadratic model to:

$$\eta_{ij} = C_{ij} - \frac{1}{2}\Big(-2\boldsymbol{z}_i^\top\boldsymbol{\gamma}_j + 2\boldsymbol{z}_i^\top\boldsymbol{D}_j\boldsymbol{z}_i\Big),$$

so that we need to integrate:

$$\int \exp\left[C_{ij} - \frac{1}{2}\left\{2\boldsymbol{z}_i^\top \boldsymbol{D}_j \boldsymbol{z}_i - 2\boldsymbol{z}_i \boldsymbol{\gamma}_j + \left(\boldsymbol{z}_i - \boldsymbol{a}_i\right)^\top \boldsymbol{A}_i^{-1}\left(\boldsymbol{z}_i - \boldsymbol{a}_i\right)\right\}\right] \det\left(\boldsymbol{A}_i\right)^{-\frac{1}{2}} d\boldsymbol{z}_i.$$

Taking out terms that are constant with respect to the latent scores, we can rewrite this to:

$$\exp\left(C_{ij} - \frac{1}{2}\boldsymbol{a}_i^\top \boldsymbol{A}_i \boldsymbol{a}_i\right) \int \exp\left\{-\frac{1}{2}\left(\boldsymbol{z}_i - v_{ij}\right)^\top \left(2\boldsymbol{D}_j + \boldsymbol{A}_i^{-1}\right)\left(\boldsymbol{z}_i - v_{ij}\right) + \frac{1}{2}\boldsymbol{v}_{ij}^\top\left(2\boldsymbol{D}_j + \boldsymbol{A}_i^{-1}\right)v_{ij}\right\} d\boldsymbol{z}_i,$$

with vector $\boldsymbol{v}_{ij}$ as:

$$\boldsymbol{z}_i^\top\left(2\boldsymbol{D}_j + \boldsymbol{A}_i^{-1}\right)\boldsymbol{v}_{ij} = \boldsymbol{z}_i^\top \boldsymbol{\gamma}_j + \boldsymbol{z}_i^\top \boldsymbol{A}_i^{-1}\boldsymbol{a}_i$$

$$\boldsymbol{v}_{ij} = \frac{1}{\boldsymbol{z}_i^\top}\left(2\boldsymbol{D}_j + \boldsymbol{A}_i^{-1}\right)^{-1}\left(\boldsymbol{z}_i^\top \boldsymbol{\gamma}_j + \boldsymbol{z}_i^\top \boldsymbol{A}_i^{-1}\boldsymbol{a}_i\right)$$

$$\boldsymbol{v}_{ij} = \left(2\boldsymbol{D}_j + \boldsymbol{A}_i^{-1}\right)^{-1}\left(\boldsymbol{\gamma}_j + \boldsymbol{A}_i^{-1}\boldsymbol{a}_i\right).$$

Concluding that $\exp\{\frac{1}{2}\boldsymbol{v}_{ij}^\top(2\boldsymbol{D}_j + \boldsymbol{A}_i^{-1})\boldsymbol{v}_{ij}\}$ is constant with respect to the latent variables too:

$$\exp\left[C_{ij} + \frac{1}{2}\left\{\boldsymbol{v}_{ij}^\top\left(2\boldsymbol{D}_j + \boldsymbol{A}_i^{-1}\right)\boldsymbol{v}_{ij} - \boldsymbol{a}_i^\top \boldsymbol{A}_i \boldsymbol{a}_i\right\}\right] \int \exp\left\{-\frac{1}{2}\left(\boldsymbol{z}_i - \boldsymbol{v}_{ij}\right)^\top \left(2\boldsymbol{D}_j + \boldsymbol{A}_i^{-1}\right)\left(\boldsymbol{z}_i - \boldsymbol{v}_{ij}\right)\right\} d\boldsymbol{z}_i,$$

which leads us to conclude that the solution comes from the normalizing constant for a normal distribution with mean $\boldsymbol{v}_{ij}$ and covariance $(2\boldsymbol{D}_j + \boldsymbol{A}_i^{-1})^{-1}$ as constant:

$$\int \exp\left\{-\frac{1}{2}\left(\boldsymbol{z}_i - \boldsymbol{v}_{ij}\right)^\top \left(2\boldsymbol{D}_j + \boldsymbol{A}_i^{-1}\right)\left(\boldsymbol{z}_i - \boldsymbol{v}_{ij}\right)\right\} d\boldsymbol{z}_i = 2\pi^{-\frac{n}{2}} \det\left(2\boldsymbol{D}_j + \boldsymbol{A}_i^{-1}\right)^{-\frac{1}{2}} \propto \det\left(2\boldsymbol{D}_j + \boldsymbol{A}_i^{-1}\right)^{-\frac{1}{2}}.$$

This results in the final solution for the expectation:

$$\mathrm{E}\left\{\exp\left(\eta_{ij}\right)\right\} = \exp\left[C_{ij} + \frac{1}{2}\left\{\boldsymbol{v}_{ij}^\top\left(2\boldsymbol{D}_j + \boldsymbol{A}_i^{-1}\right)\boldsymbol{v}_{ij} - \boldsymbol{a}_i^\top \boldsymbol{A}_i^{-1} \boldsymbol{a}_i\right\}\right] \det\left(2\boldsymbol{D}_j + \boldsymbol{A}_i^{-1}\right)^{-\frac{1}{2}} \det\left(\boldsymbol{A}_i\right)^{-\frac{1}{2}}.$$

Thus, the Poisson VA log-likelihood for the quadratic model is:

$$\log \mathcal{L}_{VA}(\boldsymbol{\Theta}, \boldsymbol{\xi}) = \sum_{i=1}^{n}\sum_{j=1}^{p}\Bigg[y_{ij}\Bigg\{C_{ij} + \boldsymbol{a}_i^\top \boldsymbol{\gamma}_j - \boldsymbol{a}_i^\top \boldsymbol{D}_j \boldsymbol{a}_i - \text{tr}\Bigg(\boldsymbol{D}_j \boldsymbol{A}_i\Bigg)\Bigg\}$$
$$- \exp\Bigg\{C_{ij} + \frac{1}{2}\Bigg((\boldsymbol{\gamma}_j + \boldsymbol{A}_i^{-1}\boldsymbol{a}_i)^\top (2\boldsymbol{D}_j + \boldsymbol{A}_i^{-1})^{-1}(\boldsymbol{\gamma}_j + \boldsymbol{A}_i^{-1}\boldsymbol{a}_i) \quad (21)$$
$$-\boldsymbol{a}_i^\top \boldsymbol{A}_i^{-1}\boldsymbol{a}_i\Bigg\}\det\Bigg\{2\boldsymbol{D}_j + \boldsymbol{A}_i^{-1}\Bigg\}^{-\frac{1}{2}}\det\Bigg\{\boldsymbol{A}_i\Bigg\}^{-\frac{1}{2}}\Bigg] + \frac{1}{2}\sum_{i=1}^{n}\Bigg\{\log\det\Bigg(\boldsymbol{A}_i\Bigg) - \text{tr}\Bigg(\boldsymbol{A}_i\Bigg) - \boldsymbol{a}_i^\top \boldsymbol{a}_i\Bigg\}.$$

#### Negative-Binomial: overdispersed counted responses

We model overdispersed counts using a Poisson-gamma compound distribution with a log-link function. One option is $y_{ij} \sim Pois(\mu_{ij}\nu_{ij})$ with overdispersion parameter $\nu_{ij} \sim \Gamma(\phi_j, \phi_j)$. This gives $\text{E}\{y_{ij}\} = \mu_{ij}$ and $\text{var}(y_{ij}) = \mu_{ij} + \mu_{ij}/\phi_j$. However, this model is unidentifiable since it can be rewritten as:

$$\eta_{ij} = C_{ij} + \boldsymbol{z}_i^\top \boldsymbol{\gamma}_j - \boldsymbol{z}_i^\top \boldsymbol{D}_j \boldsymbol{z}_i, + \log\{\nu_{ij}\}$$

where the two last terms are both indexed per site and species, and constrained to be negative only. This proof serves to demonstrate that the quadratic term of the model acts as an overdispersion term for counted data. Alternatively, from this it can be concluded that, a linear GLLVM with negative-binomial distribution can be understood as modeling tolerances that are different for species, though equal for latent variables, with the overdispersion parameter. In practice, the quadratic GLLVM should only require a negative-binomial distribution for extremely overdispersed count data, i.e. when there is overdispersion for species that also have a narrow niche. Thus, for cases of extreme overdispersion, we implement the same negative-binomial parameterization as was used in Hui et al. (2017): $y_{ij} \sim Pois(\nu_{ij})$ where $\nu_{ij} \sim \Gamma(\phi_j, \phi_j/\mu_{ij})$. This parameterization has the advantage of being identifiable as it cannot be reparameterized in the same way as above, and has a quadratic mean-variance relationship.

This parameterization requires integration of $\text{E}\{\exp(-\eta)\}$, for which the solution is similar to $\text{E}\{\exp(\eta)\}$ in the case of Poisson responses above. However, the calculation here includes the terms $(\boldsymbol{A}_i - 2\boldsymbol{D}_j)^{-1}$, which we assume to be positive semi-definite or negative semi-definite. This assumption only fails when the resulting matrix is singular, which in practice happens when the off-diagonals of $\boldsymbol{A}_i$ are zero, and the diagonals of the matrices match.

The log-likelihood of negative-ninomial distributed responses is:

$$\mathcal{L}(\boldsymbol{\Theta}) = \sum_{i=1}^{n}\sum_{j=1}^{p}\Bigg\{\Bigg(y_{ij}+\phi_j-1\Bigg)\log\Bigg(\nu_{ij}\Bigg)-\Bigg(1+\frac{\phi_j}{\mu_{ij}}\Bigg)\nu_{ij}-\phi_j\eta_{ij}+\phi_j\log\Bigg(\phi_j\Bigg)-\log\Gamma\Bigg(\phi_j\Bigg)-\log\Bigg(y_{ij}!\Bigg)\Bigg\}-\frac{1}{2}\sum_{i=1}^{n}\boldsymbol{z}_i^\top\boldsymbol{z}_i, \quad (22)$$

where terms constant as a function of the parameters have been omitted. The optimal variational distribution for the latent variable $\nu_{ij}$ is:

$$
\begin{aligned}
\log\{\mathrm{q}(\nu_{ij})\} &\propto \mathrm{E}(\mathcal{L}(\boldsymbol{\Theta})) \\
&\propto (y_{ij}+\phi_j-1)\log(\nu_{ij}) - [1+\phi_j \mathrm{E}\{\exp(-\eta_{ij})\}]\nu_{ij} \\
&\propto \left(y_{ij}+\phi_j-1\right)\log\left(\nu_{ij}\right) - \Bigg[1+\phi_j \exp\Bigg\{-C_{ij} \\
&+\frac{1}{2}\Bigg((-\boldsymbol{\gamma}_j+\boldsymbol{A}_i^{-1}\boldsymbol{a}_i)^\top(\boldsymbol{A}_i^{-1}-2\boldsymbol{D}_j)^{-1}(-\boldsymbol{\gamma}_j+\boldsymbol{A}_i^{-1}\boldsymbol{a}_i)-\boldsymbol{a}_i^\top\boldsymbol{A}_i^{-1}\boldsymbol{a}_i\Bigg)\Bigg\}\det\left(\boldsymbol{A}_i^{-1}-2\boldsymbol{D}_j\right)^{-\frac{1}{2}}\det\left(\boldsymbol{A}_i\right)^{-\frac{1}{2}}\Bigg]\nu_{ij},
\end{aligned}
$$

where terms constant as a function of the parameters have been omitted. Note, that the term $\det(\boldsymbol{A}_i^{-1}-2\boldsymbol{D}_j)^{-\frac{1}{2}}$ needs to be calculated as $\boldsymbol{B}_{ij}=\boldsymbol{A}_i^{-1}-2\boldsymbol{D}_j$ with $\boldsymbol{L}_{ij}=\mathrm{chol}(\boldsymbol{B}_{ij})$, so that $\det(\boldsymbol{A}_i^{-1}-2\boldsymbol{D}_j)^{-\frac{1}{2}}=\det(\boldsymbol{L}_{ij})^{-1}$, where $\mathrm{chol}(\cdot)$ represents the cholesky factorization. This prevents having to potentially calculate the (undefined) square root of a negative determinant.

From this we conclude that $\mathrm{q}(v_{ij})\sim\Gamma(y_{ij}+\phi_j,\zeta_{ij})$, with:

$$
\zeta_{ij}=1+\phi_j\exp\Bigg[-C_{ij}+\frac{1}{2}\Bigg\{\left(-\boldsymbol{\gamma}_j+\boldsymbol{A}_i^{-1}\boldsymbol{a}_i\right)^\top\boldsymbol{B}^{-1}\left(-\boldsymbol{\gamma}_j+\boldsymbol{A}_i^{-1}\boldsymbol{a}_i\right)-\boldsymbol{a}_i^\top\boldsymbol{A}_i^{-1}\boldsymbol{a}_i\Bigg\}\Bigg]\det\left(\boldsymbol{L}_{ij}\right)^{-1}\det\left(\boldsymbol{A}_i\right)^{-\frac{1}{2}}. \tag{23}
$$

The VA log-likelihood is derived by taking expectations over the components of equation (13):

$$
\begin{aligned}
\mathrm{E}[\log\{f(y_{ij}|\boldsymbol{z}_i,\boldsymbol{\Theta})h(\boldsymbol{z}_i)\}] &= \sum_{i=1}^{n}\sum_{j=1}^{p}\Bigg[\Big\{y_{ij}+\phi_j-1\Big\}\Big\{\psi\Big(y_{ij}+\phi_j\Big)-\log\Big(\zeta_{ij}+\phi_j\Big)\Big\}-\Big\{y_{ij}+\phi_j\Big\}-\phi_j\tilde{\eta}_{ij} \\
&\quad +\phi_j\log\Big\{\phi_j\Big\}-\log\Gamma\Big\{\phi_j\Big\}\Bigg]-\frac{1}{2}\sum_{i=1}^{n}\Big\{\mathrm{tr}\Big(\boldsymbol{A}_i\Big)+\boldsymbol{a}_i^\top\boldsymbol{a}_i\Big\} \\
\mathrm{E}[\log\{\mathrm{q}(\nu_{ij})\}] &= \sum_{i=1}^{n}\sum_{j=1}^{p}\Bigg[\Big\{y_{ij}+\phi_j-1\Big\}\Big\{\psi\Big(y_{ij}+\phi_j\Big)-\log\Big(\zeta_{ij}+\phi_j\Big)\Big\}-\Big(y_{ij}+\phi_j\Big)+\Big(y_{ij}+\phi_j\Big)\log\Big(\zeta_{ij}+\phi_j\Big) \\
&\quad -\log\Gamma\Big(y_{ij}+\phi_j\Big)\Bigg] \\
\mathrm{E}(\log\{\mathrm{q}(\boldsymbol{z}_i|\boldsymbol{a}_i,\boldsymbol{A}_i)\}) &= -\frac{1}{2}\sum_{i=1}^{n}\log\det(2e\pi\boldsymbol{A}_i).
\end{aligned}
$$

Thus, the negative-binomial VA log-likelihood for the quadratic model is:

$$\mathcal{L}_{VA}(\mathbf{\Theta}, \boldsymbol{\xi}) = \sum_{i=1}^{n}\sum_{j=1}^{p}\left\{-\phi_j\tilde{\eta}_{ij} - \left(y_{ij} + \phi_j\right)\log\left(\zeta_{ij}\right) + \log\Gamma\left(y_{ij}+\phi_j\right) + \phi_j\log\left(\phi_j\right) - \log\Gamma\left(\phi_j\right)\right\}$$
$$+\frac{1}{2}\sum_{i=1}^{n}\left\{\log\det\left(\boldsymbol{A}_i\right) - \mathrm{tr}\left(\boldsymbol{A}_i\right) - \boldsymbol{a}_i^\top\boldsymbol{a}_i\right\}. \quad (24)$$

Note, that this log-likelihood can be re-arranged similarly as in Hui et al. (2017), by replacing $\frac{1}{2}\boldsymbol{\gamma}_j^\top\boldsymbol{A}_i\boldsymbol{\gamma}_j$ with $\exp[\boldsymbol{a}_{\boldsymbol{i}}^\top\boldsymbol{\gamma}_j - \boldsymbol{a}_i\boldsymbol{D}_j\boldsymbol{a}_i - \mathrm{tr}(\boldsymbol{D}_j\boldsymbol{A}_i) + \frac{1}{2}\{(-\boldsymbol{\gamma}_j + \boldsymbol{A}^{-1}\boldsymbol{a}_i)^\top\boldsymbol{B}_{ij}^{-1}(-\boldsymbol{\gamma}_j + \boldsymbol{A}^{-1}\boldsymbol{a}_i) - \boldsymbol{a}_i^\top\boldsymbol{A}_i\boldsymbol{a}_i\}]\det(\boldsymbol{L}_{ij})^{-1}\det(\boldsymbol{A}_i)^{-0.5}$.

### Gamma responses: positive continuous

The log-likelihood for gamma distributed responses with a log-link function, shape $\frac{1}{\phi_j}$ and scale $\mu_{ij}\phi_j$ is:

$$\mathcal{L}(\mathbf{\Theta}) = \sum_{i=1}^{n}\sum_{j=1}^{p}\left[-\frac{1}{\phi_j}\left\{\eta_{ij} + \log\phi_j - \log\left(y_{ij}\right)\right\} - y_{ij}\frac{1}{\phi_j}\exp\left\{-\eta_{ij}\right\} - \log\left\{y_{ij}\right\} - \log\Gamma\left\{\frac{1}{\phi_j}\right\}\right] - \frac{1}{2}\sum_{i=1}^{n}\boldsymbol{z}_i^\top\boldsymbol{z}_i, \quad (25)$$

where terms constant as a function of the parameters have been omitted. The VA log-likelihood is derived by taking expectations over the components of equation (4) with respect to the latent variables:

$$\mathrm{E}[\log\{f(y_{ij}|\boldsymbol{z}_i,\mathbf{\Theta})h(\boldsymbol{z}_i)\}] = \sum_{i=1}^{n}\sum_{j=1}^{p}\left[-\frac{1}{\phi_j}\left\{\mathrm{E}\left(\eta_{ij}\right) + \log\phi_j - \log\left(y_{ij}\right)\right\} - y_{ij}\frac{1}{\phi_j}\mathrm{E}\left\{\exp\left(-\eta_{ij}\right)\right\}$$
$$-\log\left(y_{ij}\right) - \log\Gamma\left\{\frac{1}{\phi_j}\right\}\right] - \frac{1}{2}\sum_{i=1}^{n}\left\{\mathrm{tr}\left(\boldsymbol{A}_i\right) + \boldsymbol{a}_i^\top\boldsymbol{a}_i\right\}$$

$$\mathrm{E}[\log\{\mathrm{q}(\boldsymbol{z}_i|\boldsymbol{a}_i,\boldsymbol{A}_i)\}] = -\frac{1}{2}\sum_{i=1}^{n}\log\det(2e\pi\boldsymbol{A}_i).$$

The result for $\mathrm{E}(\eta_{ij})$ has been given above for all distributions, and the result for $\mathrm{E}\{\exp(-\eta_{ij})\}$ in the derivation of the VA log-likelihoood for the negative-binomial distribution. Thus, the gamma VA log-likelihood for the quadratic model is:

$$
\begin{aligned}
\mathcal{L}(\boldsymbol{\Theta}, \boldsymbol{\xi}) = \sum_{i=1}^{n}\sum_{j=1}^{p}\Bigg[&-\frac{1}{\phi_j}\Big\{\eta_{ij} + \log\Big(\phi_j\Big) - \log\Big(y_{ij}\Big)\Big\} - y_{ij}\frac{1}{\phi_j}\exp\Big\{-C_{ij} \\
&+ \frac{1}{2}\Big((-\boldsymbol{\gamma}_j + \boldsymbol{A}_i^{-1}\boldsymbol{a}_i)^\top \boldsymbol{B}_{ij}^{-1}(-\boldsymbol{\gamma}_j + \boldsymbol{A}_i^{-1}\boldsymbol{a}_i) - \boldsymbol{a}_i^\top \boldsymbol{A}_i^{-1}\boldsymbol{a}_i\Big)\Big\} \det\Big\{\boldsymbol{L}_{ij}\Big\}^{-1} \det\Big\{\boldsymbol{A}_i\Big\}^{-\frac{1}{2}} - \log\Big\{y_{ij}\Big\} \\
&- \log\Gamma\Big\{\frac{1}{\phi_j}\Big\}\Bigg] + \frac{1}{2}\sum_{i=1}^{n}\log\det\Big(\boldsymbol{A}_i\Big) - \mathrm{tr}\Big(\boldsymbol{A}_i\Big) - \boldsymbol{a}_i^\top\boldsymbol{a}_i,
\end{aligned} \tag{27}
$$

with $\boldsymbol{B}_{ij} = \boldsymbol{A}_i^{-1} - 2\boldsymbol{D}_j$ and $\boldsymbol{L}_{ij} = \mathrm{chol}(\boldsymbol{B}_{ij})$.

#### Approximate standard errors

As noted by Hui et al. (2017), approximate asymptotic standard errors for parameters can be retrieved from the Hessian matrix of the VA log-likelihood. Here, we provide a brief overview of the Delta method, that can be used for the calculation of additional approximate standard errors for the species optima $\boldsymbol{u}_j$, tolerances $\boldsymbol{t}_j$, species maxima $\boldsymbol{c}_j$, and gradient length, per latent variable. The multivariate Delta method states, that approximate standard errors can be calculated for a parameter $\theta$ that is a function $f(\cdot)$ of a set of other parameters, by the gradient of the function with respect to the original parameters $\nabla \mathrm{f}(\cdot)$, and the variance-covariance matrix $\boldsymbol{\Sigma}$ of the original parameters. Assuming independence between species and latent variables, the variance of the species optima, maxima, or tolerances, is given by:

$$
\mathrm{Var}\{\mathrm{f}(\theta)\} \approx \nabla\mathrm{f}(\gamma_{jq}, D_{jqq})^\top \boldsymbol{\Sigma} \nabla\mathrm{f}(\gamma_{jq}, D_{jqq}),
$$

where $\theta$ is the optimum, maximum, or tolerance of a species response curve for a latent variable, or the gradient length, and $\nabla\mathrm{f}(\gamma_{jq}, D_{jqq})$ is the first derivative of $\theta$ with respect to the separate parameters $\gamma_{jq}$ and $D_{jqq}$. Tolerances and gradient lengths are only a function of the quadratic coefficient(s), not of the linear coefficients.

### Appendix S4: Fitting process

In this appendix, details on stabilizing the fitting on quadratic GLLVMs are included. We used Template Model Builder (TMB; Kristensen et al., 2016) to retrieve analytical derivatives and more smoothly fit the models.

#### Fitting

Even with the use of VA, estimation of GLLVMs is particularly challenging due to multimodality of the log-likelihood function. This often results in numerical optimization techniques getting stuck in local minima. We largely adopt the approach developed by Niku et al. (2019), who found that fitting GLLVMs with diagonal VA covariance matrices before fitting the model with unstructured VA covariance matrices, tended to stabilize the estimation for the linear GLLVM (Hui et al., 2017). So, to overcome this problem, we recommend to first fit a species-common tolerances model with diagonal VA covariance matrix, i.e. a model where $\boldsymbol{D}_j = \boldsymbol{D}$, after which the solution is used as initial values to fit a species-common tolerances model with full VA covariance matrix, of which the solution is used as the initial values to fit the species-specific quadratic model. In practice, we found this procedure to stabilize fitting of the quadratic GLLVM. Alternatively, the quadratic GLLVM can be fitted with diagonal or full VA covariance matrix immediately, thus providing different fitting algorithms. Initial values for the numerical optimization are generated following the same procedure implemented in Niku et al. (2019), with the most basic option to start the optimization with a combination of zeros for the coefficients and ones for the latent variables, or with randomly generated initial values, or with initial values generated by: 1) fitting a multivariate Generalized Linear Model, 2) retrieving the Dunn-Smyth residuals (Dunn and Smyth, 1996) of the model, and 3) performing factor analysis on those residuals. It is also possible to use the solution of a linear GLLVM as initial values, though this usually performed either similar or worse than the approach for initial values described by Niku et al. (2019). When using the solution of a linear GLLVM as initial values, the options for the linear GLLVM can be tweaked in addition to tweaking the algorithm for the quadratic GLLVM (e.g. by running it multiple times with different sets of initial values), resulting in an even larger number of potential fitting algorithms. In general, initial values that are closer to the final solution serve to further reduce the possibility for the optimizer to get stuck in local minima. Fitting the model with different sets of initial values can help to assess if is has properly converged.

Estimation of the quadratic coefficients can be started at a small constant, or on a solution that is maximized conditionally on the initial values. Maximizing conditionally on the initial values further reduces the possibility for the optimizer to get stuck in local minima, in scenarios where the initial values are close to the optimal solution. This approach works especially well in combination with first fitting the species-common tolerances model. It is possible to assess if a model converged to the solution that optimally maximizes the VA log-likelihood by examining the analytical gradient, in combination with the standard errors. The gradient should be close to zero if the model has converged. In our experience, the standard errors tend to be larger when a model has not converged.

#### Regularisation

The quadratic model can be further extended with an additional species-common component, by changing the quadratic term to $\boldsymbol{z}_i^\top (\boldsymbol{D}_j + \boldsymbol{G}) \boldsymbol{z}_i$, where $\boldsymbol{G}$ is again a positive-definite diagonal matrix. Including a species-common component allows to make optimal use of the data, as parameters that are equal for all species can potentially be estimated with a higher degree of certainty than species-specific components $\boldsymbol{G}$. In this case, $\boldsymbol{D}_j$ accounts for species-specific deviation from the species-common components. The species-common components can be interpreted as an average measure of tolerance, as gradient length, or as compositional turnover. This parameterization is unidentifiable, thus constraints for the species-specific component are required. For example, penalized likelihood methods could be used to enforce identifiability (Tibshirani, 1996). This parameterization provides a hybrid form of the species-common and species-specific tolerances models, as with a large penalty it will fit the species-common tolerances model, and with a small or no penalty the species-specific tolerances model. This can additionally serve to further stabilize the fitting process (as discussed above and in Yee, 2004). Alternatively, the elements of $\boldsymbol{D}_j$ can be treated as a random effect, independent for species $p$ and latent variables $d$, following a half-normal distribution, although we leave this as an avenue of future research.

### Appendix S5: Extra material for simulations quadratic GLLVM

#### Figure S1: Initial values

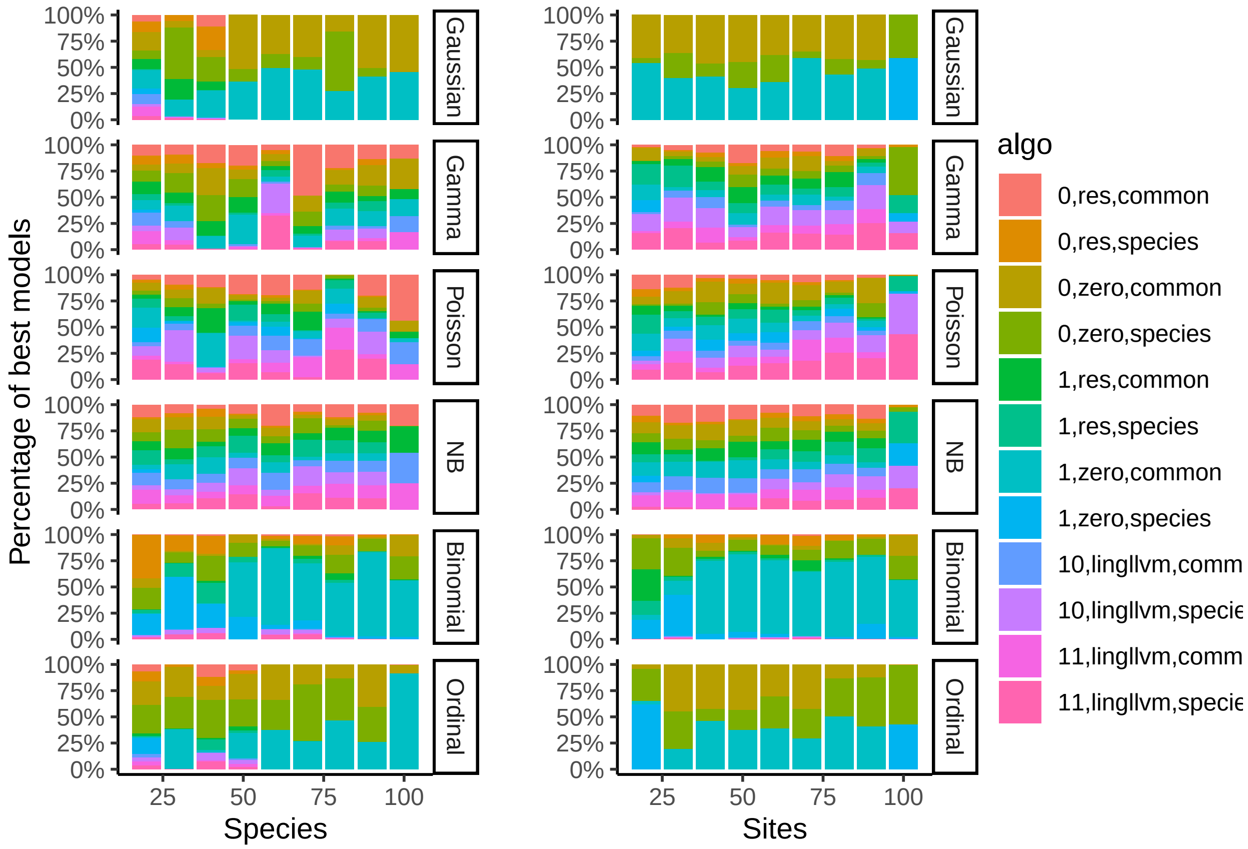


Figure 1: Starting values from best models of the simulated datasets. The legend indicates settings of the model, with in order: model with diagonal VA matrix fitted first (0,1), starting values used (zero, sensibly generated (res), or linear GLLVM), and whether the first a common tolerances was fit (common) or not (species) before the full model.

#### Figure S2: MAE with all species optima

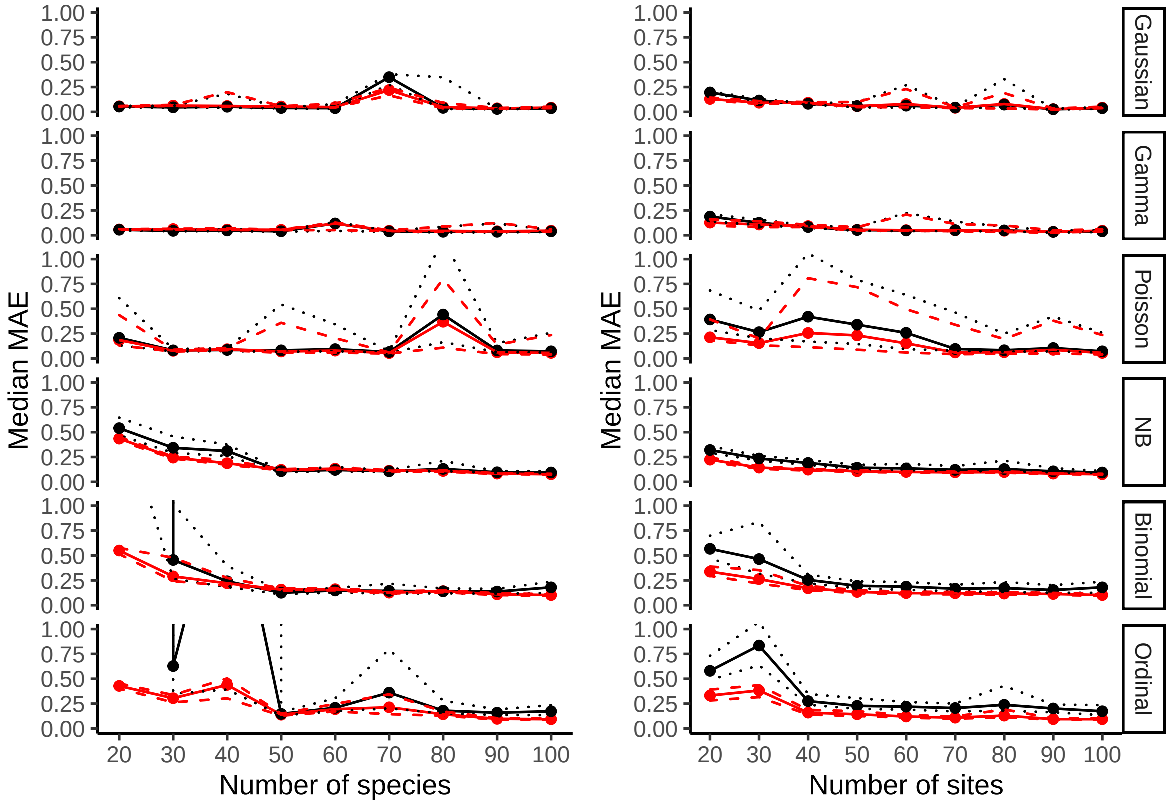


Figure 2: Simulation results for the quadratic GLLVM per distribution, only optima that could be estimated are included (larger than 10 or smaller than -10 were excluded). The left column shows simulations where the number of sites was kept constant at 100, the right the same for the number of species. Twelve models were fit to each simulated dataset from which the best was kept as the final model. The figure includes the median MAE, shown as dots connected by solid lines, for species optima (black) and latent variables (red), with the first and third quantiles represented as dotted (optima) and dashed (latent variables) lines.

### Appendix S5: Results for example implementation of the hunting spider dataset

| | latent variables | df | LL | AIC | delta AIC | AIC weight |
|---|---|---|---|---|---|---|
| 4 | 4 | 55.00 | -762.55 | 1415.09 | 0.00 | 0.99 |
| 5 | 5 | 63.00 | -761.63 | 1425.27 | 10.17 | 0.01 |
| 3 | 3 | 46.00 | -787.74 | 1439.90 | 24.80 | 0.00 |
| 2 | 2 | 36.00 | -837.99 | 1451.98 | 36.89 | 0.00 |
| 1 | 1 | 25.00 | -1195.47 | 3090.94 | 1675.84 | 0.00 |

Table S1: Model selection results for linear GLLVM fit to the hunting spider dataset, to find the number of latent variables.

| | tolerances | df | LL | AIC | delta AIC | AIC weight |
|---|---|---|---|---|---|---|
| 3 | Species-specific | 59 | -729.09 | 1354.93 | 1.00 | 1.00 |
| 2 | Equal | 37 | -827.71 | 1448.22 | 0.00 | 0.00 |
| 1 | Species-common | 36 | -837.99 | 1451.98 | 0.00 | 0.00 |

Table S2: Model selection results of species tolerances for the hunting spider dataset.

| | latent variables | df | LL | AIC | delta AIC | AIC weight |
|---|---|---|---|---|---|---|
| 3 | 3 | 81.00 | -674.49 | 1264.98 | 0.00 | 0.95 |
| 4 | 4 | 102.00 | -673.43 | 1270.71 | 5.72 | 0.05 |
| 5 | 5 | 122.00 | -693.10 | 1314.28 | 49.30 | 0.00 |
| 2 | 2 | 59.00 | -729.09 | 1354.93 | 89.95 | 0.00 |
| 1 | 1 | 36.00 | -1278.96 | 2333.92 | 1068.94 | 0.00 |

Table S3: Model selection results for the quadratic GLLVM fit to the hunting spider dataset, to find the number of latent variables.

| | diag.iter | starting.val | start.struc | n.init | jitter.var | df | LL | AIC | delta AIC | AIC weight | SE |
|---|---|---|---|---|---|---|---|---|---|---|---|
| 3 | yes | zero | species | 0.00 | no | 81 | -670.67 | 1257.34 | 0.00 | 0.42 | yes |
| 9 | no | linear GLLVM | species | 3.00 | yes | 81 | -671.59 | 1259.19 | 1.85 | 0.17 | no |
| 7 | yes | res | species | 5.00 | no | 81 | -671.66 | 1259.33 | 1.99 | 0.16 | yes |
| 8 | yes | res | common | 5.00 | no | 81 | -672.29 | 1260.58 | 3.25 | 0.08 | yes |
| 12 | yes | linear GLLVM | common | 3.00 | yes | 81 | -672.29 | 1260.58 | 3.25 | 0.08 | no |
| 6 | no | res | common | 5.00 | no | 81 | -672.43 | 1260.87 | 3.53 | 0.07 | no |
| 2 | no | zero | common | 0.00 | no | 81 | -674.49 | 1264.98 | 7.65 | 0.01 | yes |
| 5 | no | res | species | 5.00 | no | 81 | -675.55 | 1267.10 | 9.77 | 0.00 | yes |
| 4 | yes | zero | common | 0.00 | no | 81 | -675.98 | 1267.96 | 10.62 | 0.00 | yes |
| 10 | no | linear GLLVM | common | 3.00 | yes | 81 | -680.21 | 1276.41 | 19.08 | 0.00 | no |
| 11 | yes | linear GLLVM | species | 3.00 | yes | 81 | -685.01 | 1286.02 | 28.69 | 0.00 | yes |
| 1 | no | zero | species | 0.00 | no | 81 | -685.01 | 1286.03 | 28.69 | 0.00 | yes |

Table S4: Model-selection results for the final quadratic GLLVM for the hunting spider dataset. The columns diag.iter, start.struc, n.init and jitter.var correspond to options for the fitting algorithm. Diag.iter first fits a model with diagonal covariance matrix before the full model. Start.struc first fits a model with species-common tolerances before the full model. N.init performs multiple models runs, and picks the best solution by the highest log-likelihood (LL). Jitter.var adds a small amount of gaussian noise (as much as put in for the variance, here 0.2), to generate quasi-random starting values. The SE column indicates whether it was possible to calculate a matrix of standard errors (if not, the model has not converged)

#### Figure S3: Plot of coefficients for elevation, from the Swiss alpine plants dataset

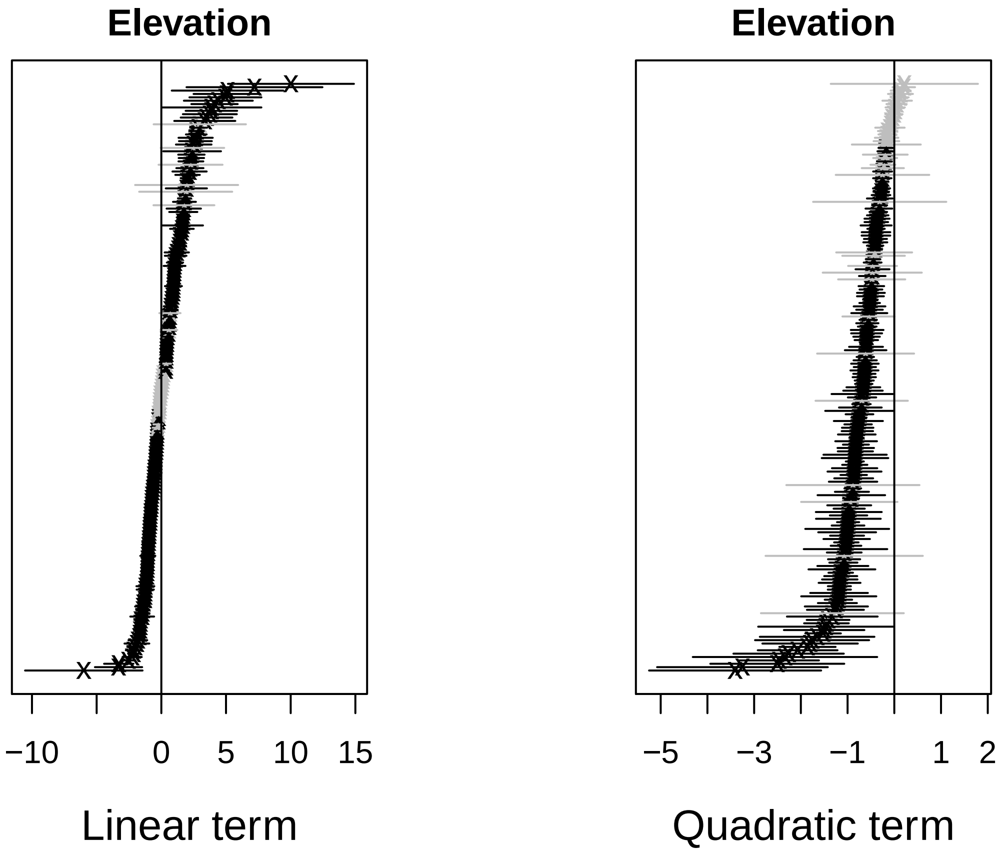

### References


P. K. Dunn and G. K. Smyth. Randomized Quantile Residuals. *Journal of Computational and Graphical Statistics*, 5(3):236–244, 1996. ISSN 1061-8600. doi: 10.2307/1390802.

F. K. C. Hui, D. I. Warton, J. T. Ormerod, V. Haapaniemi, and S. Taskinen. Variational Approximations for Generalized Linear Latent Variable Models. *Journal of Computational and Graphical Statistics*, 26(1): 35–43, Jan. 2017. ISSN 1061-8600. doi: 10.1080/10618600.2016.1164708.

K. Kristensen, A. Nielsen, C. W. Berg, H. Skaug, and B. Bell. TMB: Automatic Differentiation and Laplace Approximation. *Journal of Statistical Software*, 70(5), 2016. ISSN 1548-7660. doi: 10.18637/jss.v070.i05.

J. Niku, W. Brooks, R. Herliansyah, F. K. C. Hui, S. Taskinen, and D. I. Warton. Efficient estimation of generalized linear latent variable models. *PLOS ONE*, 14(5):e0216129, May 2019. ISSN 1932-6203. doi: 10.1371/journal.pone.0216129.

J. T. Ormerod and M. P. Wand. Explaining Variational Approximations. *The American Statistician*, 64(2): 140–153, May 2010. ISSN 0003-1305. doi: 10.1198/tast.2010.09058.

R. Tibshirani. Regression Shrinkage and Selection Via the Lasso. *Journal of the Royal Statistical Society: Series B (Methodological)*, 58(1):267–288, 1996. ISSN 2517-6161. doi: 10.1111/j.2517-6161.1996.tb02080.x.

T. W. Yee. A New Technique for Maximum-Likelihood Canonical Gaussian Ordination. *Ecological Monographs*, 74(4):685–701, 2004. ISSN 1557-7015. doi: 10.1890/03-0078.
